# Disentangling the regulatory response of *Agrobacterium tumefaciens* CHLDO to glyphosate for engineering whole-cell phosphonate biosensors

**DOI:** 10.1101/2024.07.19.604230

**Authors:** Fiorella Masotti, Nicolas Krink, Nicolas Lencina, Natalia Gottig, Jorgelina Ottado, Pablo I. Nikel

## Abstract

Phosphonates (PHTs), organic compounds with a stable C—P bond, are widely distributed in nature. Glyphosate (GP), a synthetic PHT, is extensively used in agriculture and has been linked to various human health issues and environmental damage. Given the prevalence of GP, developing cost-effective, on-site methods for GP detection is key for assessing pollution and reducing exposure risks. We adopted *Agrobacterium tumefaciens* CHLDO, a natural GP degrader, as the source of genetic parts for constructing PHT biosensors. In this species, the *phn* gene cluster, encoding the C—P lyase pathway, is regulated by the PhnF transcriptional repressor and is part of the Pho regulon. We selected the *phnG* promoter, which displays a dose-dependent response to GP, to build a set of whole-cell biosensors. Through stepwise optimization of the transcriptional cascade, we created a biosensor capable of detecting GP in the 0.25-50 μM range in various samples, including soil and water.

## Introduction

Phosphonates (PHTs) are organic compounds containing a chemically-stable carbon—phosphorus (C— P) bond. PHTs are widespread in nature [1], where they act as a source of phosphorus for microbial growth (e.g., phosphonopyruvate and phosphonoacetate) and as natural antibiotics (e.g., fosfomycin and bialaphos) [2]. Some industrial PHTs, however, rank as xenobiotics, owing to their use as herbicides [3], e.g., *N*-(phosphonomethyl)glycine, commonly known as glyphosate (GP; C_3_H_8_NO_5_P), and the GP-derivative (aminomethyl)phosphonate (AMPA; CH_6_NO_3_P). Besides applications in agriculture and silviculture, other PHTs of environmental concern are ethyl-and phenyl-phosphonates, typically employed as insecticides; alafosfalin and phosphonomycin (i.e., bisphosphonate), used as antibiotics; and cyclic esters of aromatic bisphosphonates, which are exploited as polymeric additives in the material sector [4]. Out of these examples, PHTs used in agriculture are, by far, the most problematic as they uncontrollably enter the environment and become toxic above certain threshold concentrations [5]. GP-based herbicides (GBHs) are the most common synthetic agrochemicals, with 600,000 to 750,0000 tons produced annually [6]. GP inhibits the 5-enolpyruvylshikimate 3-phosphate synthetase within the shikimate pathway in plants [7], disrupting the synthesis of aromatic amino acids and secondary metabolites. Genetically-modified crops expressing a GP-resistant enzyme of bacterial origin (e.g., from *Agrobacterium* [8]) enabled the massive employment of GBHs, triggering a critical, global-scale evaluation of the use of GBHs and their environmental and health risks [9–11]. In Europe, GP has been detected in water sources at concentrations up to 165 μg/L [12], well above the maximal allowed levels (100 μg/L) [13].

The substantial release of PHTs in the environment prompted interest in studying and understanding the degradation and metabolic processing of these molecules by microorganisms [14,15]. Microbial GP catabolism proceeds by either of two pathways [16,17]. One of these involves the cleavage of the C—P bond by a C—P lyase [18], resulting in the formation of *N*-methylglycine (i.e., sarcosine, later assimilated *via* the endogenous one-carbon metabolism) and a phosphorus-containing moiety. The second route proceeds through breaking a carboxymethylene-nitrogen bond by an O_2_-dependent oxidase [19], leading to the formation of AMPA and glyoxylate. Glyoxylate is processed via the glyoxylate shunt, whereas AMPA is acetylated and the C—P bond is cleaved through a set of reactions that include the C—P lyase. The C—P lyase enzyme system is widespread among bacteria; *Escherichia coli*, for instance, harbors the *phn* operon that encodes functions related to PHT conversion [20] (although this species is unable to use GP as phosphorus source). The transcriptional pattern of these genes seems to be subjected to a dual regulatory mechanism, i.e., upregulation under phosphate (Pi) starvation, mediated by the PhoR/PhoB two-component signal transduction system [21,22], and PhnF-mediated repression [23]. The accepted rationale behind such dual mechanism is that expression of the *phn* operon requires Pi deprivation, and it is strictly essential when PHTs are present in the medium. Therefore, under Pi-limiting conditions and in the absence of PHTs, the *phn* genes would remain unexpressed due to PhnF-mediated repression [17].

Multiple PHT-degrading organisms have been described and characterized at both the physiological and metabolic level, including *Ensifer meliloti* [24], *Brucella anthropi* [25], and *Burkholderia pseudomallei* [26]. Recently, 62 bacterial strains were isolated from agricultural soils based on their ability to use GP as the only phosphorus source [27]. One isolate, identified as *Agrobacterium tumefaciens* CHLDO, completely degraded 180 μg/mL GP in 96 h. While the potential of these bacterial isolates for biodegradation of GP and related PHT contaminants is yet to be exploited, detecting and quantifying these molecules remain challenging. Some PHTs, e.g., GP and AMPA, share structural and physicochemical properties with amino acids, including low UV absorption, scarce autofluorescence, and limited vaporization upon heating [28]. The quantification of these compounds by gas chromatography or high-performance liquid chromatography (HPLC) coupled to UV or fluorescence detection is difficult, unless time-consuming derivatization procedures are implemented [29]. While commonplace, these techniques are carried out only in specialized laboratories that may require specific instruments operated by experts. Alternative detection methods, e.g., biosensors [30], emerge as a promising alternative for reproducible and simple quantification of PHTs both in laboratory experiments and environmental samples. Whole-cell biosensors (WCBs) have been used, among other applications, to detect pollutants in low-cost and simple setups [31]. WCBs based on transcription factors (TF) exploit at least one TF that binds the target analyte(s), TF-responsive promoters, and downstream reporters, e.g., genes encoding fluorescent proteins or luciferase [32–34]. Hence, the TF-dependent response relies on the concentration of the analyte (inducer), resulting in an output signal [35,36]. To the best of our knowledge, WCBs have not yet been utilized for the detection of PHTs.

In this study, we characterized the response of the promoters that drive the expression of the *phn* gene cluster of *A*. *tumefaciens* CHLDO to PHTs. Building on the native transcriptional response of the *phn* promoters to GP, we adopted a modular approach for the construction of a set of TF-based WCBs for detection of PHTs. The biosensor was optimized by engineering the genetic circuit controlled by the PhnF repressor. The versatility of the resulting WCB was demonstrated in three different applications, i.e., studying microbial GP biodegradation in a laboratory setup, quantification of GBHs in soil samples, and direct herbicide detection in environmental samples by the naked eye. Taken together, our results highlight the value of repurposing the native genetic circuits of environmental bacterial isolates for the sake of whole-cell biosensing.

## Results and Discussion

### In silico identification of PHT-responsive promoters in the phn gene cluster of A. tumefaciens CHLDO

*A. tumefaciens* CHLDO, isolated from agricultural soils in Argentina, degrades GBHs using the end-products as a phosphorus source [27]. Upon sequencing the genome of strain CHLDO, PHT assimilation was associated to the presence of the *phn* gene cluster, displaying a structure similar to those described for other members of the same bacterial genus. Composed of 15 individual open reading frames (ORFs), this cluster spans the *phnFGHIJKLOCDE2E1 duf1045 phnMN* genes within a 13.3-kb long DNA segment (**Fig. 1A**). These genes encode the structural proteins involved in the uptake and degradation of PHT. In addition to these functions, PhnF has been identified as a transcriptional regulator for the genes that encode both the C—P lyase complex (PhnGHIJKL) and an aminoalkylphosphonate *N*-acetyltransferase (PhnO), essential for the utilization of α-aminoalkylphosphonates. The *phnDCE1E2* ORFs encode an ABC cassette-type transport system, and *duf1045-phnMN* leads to the production of key accessory proteins needed for the activity of the C—P lyase.

**Figure 1.**
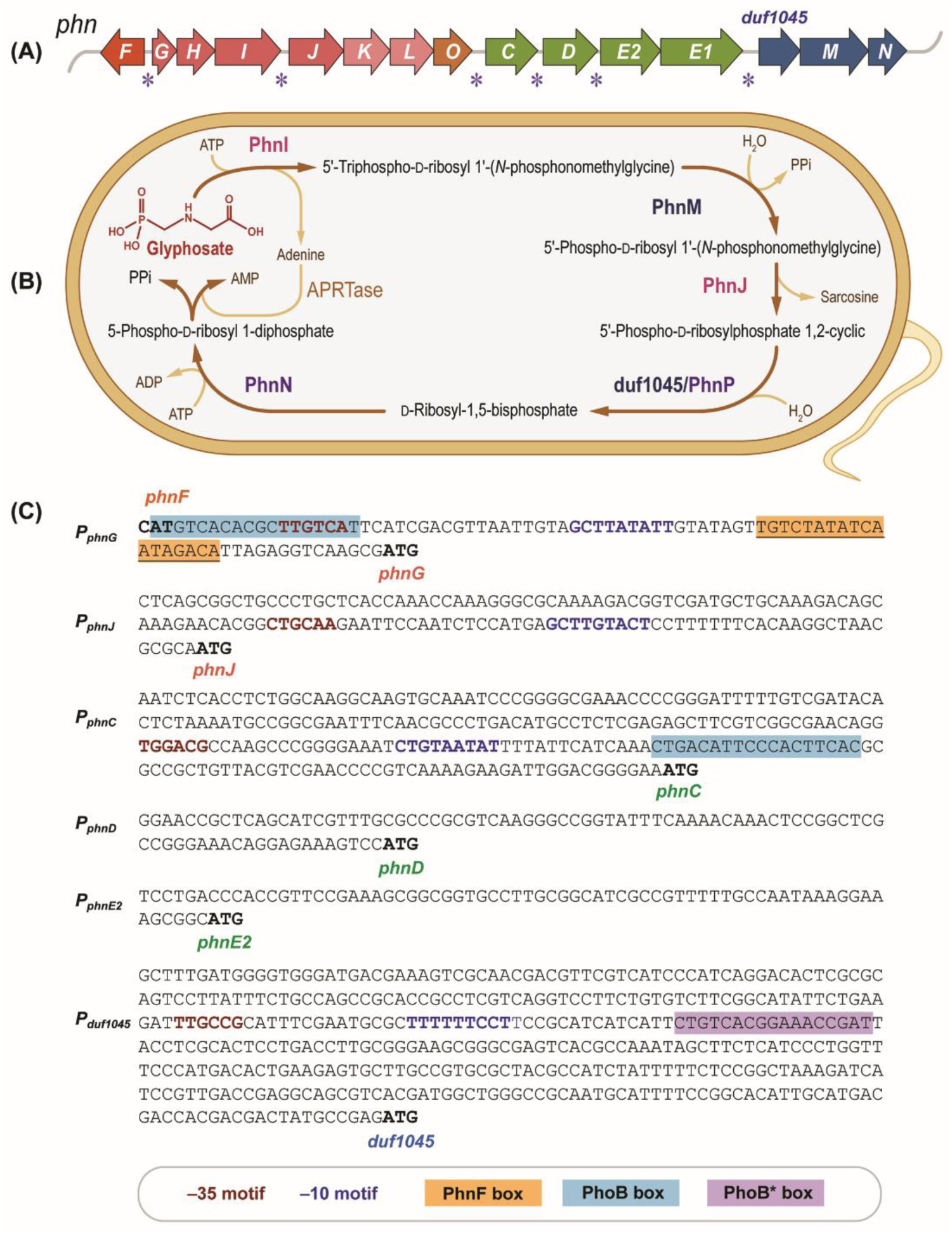
*In silico* analysis of key elements encoded in the *phn* gene cluster of *A. tumefaciens* CHLDO. **(A)** Structure of the *phn* gene cluster of *A. tumefaciens* CHLDO, encoding the carbon— phosphorus (C—P) lyase pathway. Each open reading frame is shown as an arrow; individual *phn* genes are indicated in italics. Asterisks identify putative *phn* promoters. **(B)** Reactions involved in the conversion of glyphosate [*N*-(phosphonomethyl)glycine] by the C—P lyase pathway. PhnI is a purine ribonucleoside 5′-tri/diphosphate phosphonylase, forming a complex with PhnG, PhnH, PhnJ, and PhnK. PhnM is a 5′-triphospho-α-D-ribosyl 1′-phosphonate diphosphohydrolase; PhnJ represents an *S*-adenosyl-L-methionine–dependent C—P lyase. The conversion of 5-phospho-D-ribosyl 1-diphosphate and adenine into AMP and pyrophosphate (PPi) is catalyzed by the native adenine phosphoribosyltransferase (APRTase), not encoded within the *phn* genes. **(C)** *In silico* analysis of putative promoters in the *phn* gene cluster. The *START* codons of the corresponding *phn* genes are marked in bold; the –35 and –10 sequences of the cognate promoters are identified in red and blue, respectively. Binding sites for the PhnF and PhoB transcription factors (the PhnF and PhoB box) are highlighted in orange and blue, respectively; a putative PhoB box is highlighted in purple.

Even when the responsible genes have been identified, the uptake and degradation of PHTs through the C—P lyase pathway are only partially described in *Agrobacterium*. A mechanism has been proposed for the degradation of GP by strain CHLDO through the C—P lyase pathway (**Fig. 1B**), based on the biochemical information on the processing of structurally related PHTs in *E. coli* [37,38]. This pathway releases sarcosine, with pyrophosphate (PPi) and AMP as end-products. The proposed biochemical mechanism is supported by the increased expression of *phnF*, *phnGIJKLO*, and *phnCDE2E1* when *A*. *tumefaciens* CHLDO was grown in the presence of GP as the only phosphorus source. Both *phnF* and *phnI* had 2- and 3-fold increased expression, respectively, followed by *phnJ, phnK, phnL, phnD, phnE1*, and *phnE2*, with >25-fold elevated expression. The transcription of the *phnG, phnH, phnO*, and *phnC* ORFs, on the other hand, was induced by >250-fold in the presence of GP. Interestingly, the polycistronic unit composed of *duf1045, phnM*, and *phnN* showed no significant changes in the cognate transcripts upon exposure of *A*. *tumefaciens* CHLDO to GBH [27]. These observations flagged some of the individual *phn* promoters as functional parts that can be adopted for constructing transcriptional biosensors.

We conducted an *in-silico* analysis of the *phn* cluster, focusing on the six intergenic regions between the *phn* coding sequences, i.e., *phnF-phnG*, *phnI-phnJ*, *phnO-phnC*, *phnD-phnE2*, *phnE2-phnE1*, and *phnE1-duf1045* (**Fig. 1A**). These chromosomal sequences were screened using BPROM [39] and RegPrecise [40], searching for –10 and –35 motifs and TF-binding sites. The analysis identified putative promoters upstream of the *phnG*, *phnJ*, *phnC*, *phnD*, *phnE2*, and *duf1045* ORFs, which were termed P*_phnG_,* P*_phnJ_,* P*_phnC_,* P*_phnD_*, P*_phnE2_*, and P*_duf1045_*, respectively (**Fig. 1C**). All the intergenic regions contained well defined –10 and –35 boxes, except for the 5′-unstranslated regions upstream of P*_phnD_* and P*_phnE2_*, for which these features were not evident. Binding motifs for PhnF and PhoB were likewise identified; only P*_phnG_* and P*_phnC_* contained a clearly defined PhoB box. Interestingly, a putative PhoB box sequence was also found in P*_duf1045_* (referred to as PhoB*, **Fig. 1C**). The canonical 18-nucleotide PhnF box, comprising two inverted sequences (5′-TGTCTAT-3′) separated by 4 nucleotides, was exclusively identified upstream of the P*_phnG_* promoter.

### Transcriptional response of six phn promoters upon exposure of A. tumefaciens CHLDO to GP

To investigate whether the putative P*_phnC_,* P*_phnD_,* P*_phnE2_,* P*_phnG_*, P*_phnJ_*, and P*_duf1045_* promoters are transcriptional affected by the presence of PHTs, each region was individually cloned upstream of a reporter gene encoding mScarlet, a red fluorescent protein (**Table S1** in the Supporting Information). We used Golden Gate assembly to build a first set of reporter plasmids in the versatile pL1 backbone (**Table S2** and **S3** in the Supporting Information), based on seven individual functional elements, i.e., 5’-connector, ribosome binding site (RBS), *mScarlet*, transcriptional terminator, 3’-connector, origin of replication (*oriV*), and an antibiotic (tetracycline) resistance cassette (**Fig. 2A**). These components are shared across all biosensor plasmids (**Table S1-S3** in the Supporting Information), and they were sourced from the Marburg Collection of synthetic biology parts owing to their compatibility with different microbial hosts [41]. The resulting constructs, plasmids pL1_P*_phnC_*, pL1_P*_phnD_*, pL1_P*_phnE2_*, pL1_P*_phnG_*, pL1_P*_phnJ_*, and pL1_P*_duf1045_* (**Table 1**), were introduced into *A. tumefaciens* CHLDO by electroporation. In this way, the transcriptional response of the promoters could be studied while keeping the native regulatory mechanisms unaffected. The corresponding WCB strains were termed CHLDO·P*_phnG_*, CHLDO·P*_phnJ_*, CHLDO·P*_phnC_*, CHLDO·P*_phnD_*, CHLDO·P*_phnE2_*, and CHLDO·P*_duf1045_* (**Table S4** in the Supporting Information). To establish a reference for comparisons, a control plasmid was constructed, lacking a functional promoter region (vector pL1_P*_dummy_*); this control vector was likewise transformed into strain CHLDO, giving rise to CHLDO·P*_dummy_*.

**Figure 2.**
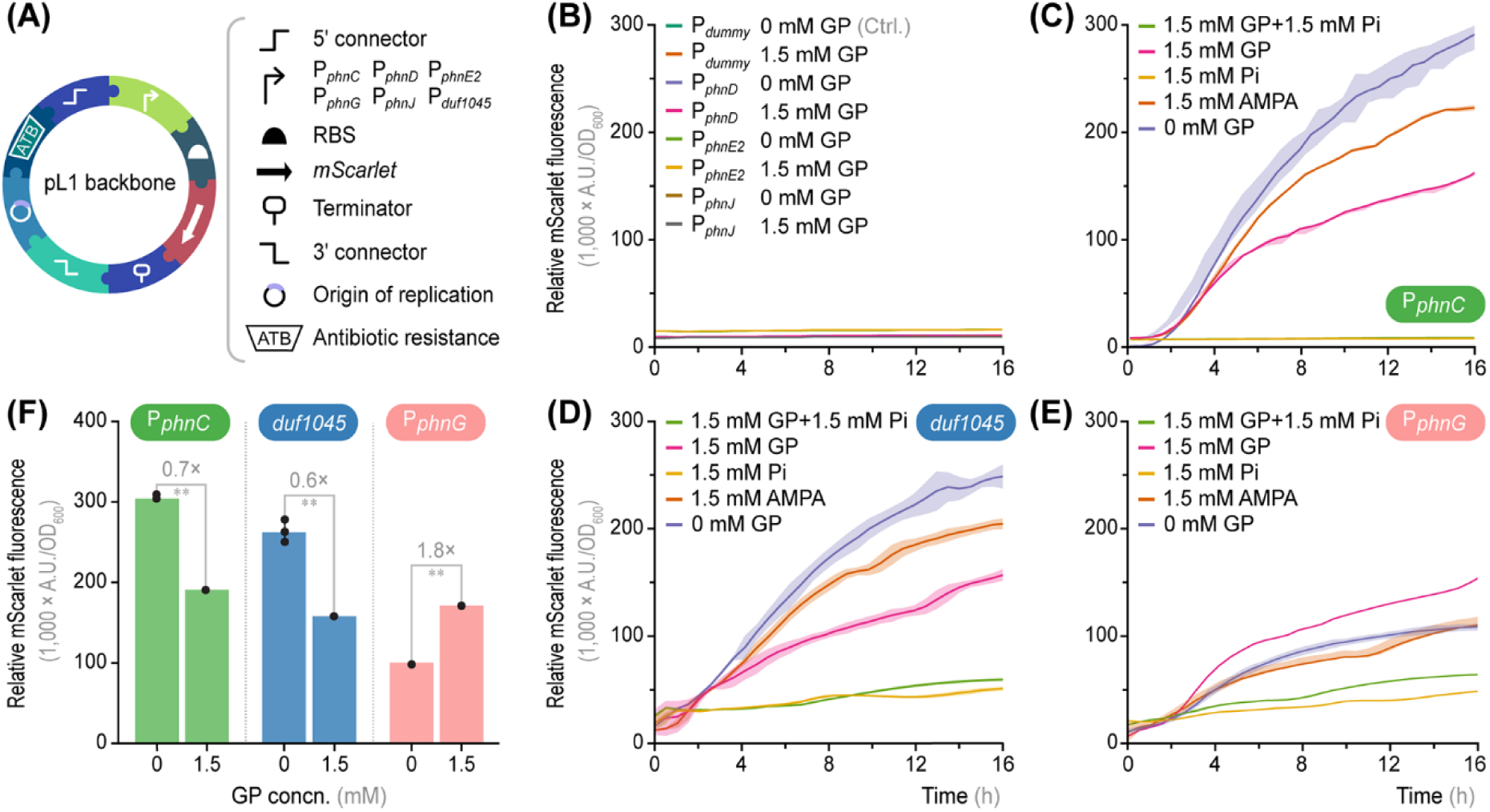
Design and test of the first set of glyphosate whole-cell biosensors. **(A)** Functional parts used to construct plasmid-borne biosensors by using different promoters of the *phn* gene cluster. Six *phn* promoters (i.e., P*_phnC_*, P*_phnD_*, P*_phnE2_*, P*_phnG_*, P*_phnJ_*, and P*_duf1045_*) were implemented in the pL1 vector backbone to drive the expression of *mScarlet* by adopting a standard format. *RBS*, ribosome binging site. Elements not drawn to scale. **(B-E)** *A*. *tumefaciens* CHLDO was individually transformed with plasmids pL1_P*_phnC_*, pL1_P*_phnC_*, pL1_P*_phnD_*, pL1_P*_phnE2_*, pL1_P*_phnG_*, pL1_P*_phnJ_*, pL1_P*_duf1045_*, and pL1_P*_dummy_* (used as a control, *ctrl*.). The output of these whole-cell biosensors (red fluorescence) was measured by incubating the cells in minimal medium with different phosphorus sources [glyphosate, GP; (aminomethyl)phosphonate, AMPA; and inorganic phosphate, Pi] for 16 h. A culture without any added phosphorus source was used as negative control; the basal mScarlet fluorescence was subtracted from the other experimental conditions. The biosensor output is expressed as arbitrary fluorescence units (*A.U.*) relative to the optical density at 600 nm (OD_600_). Shaded lines represent standard deviations for triplicate measurements from at least three independent experiments. **(F)** Maximum output for the whole-cell biosensors harboring the P*_phnC_*, P*_duf1045_*, and P*_phnG_* constructs after 16 h in the presence (1.5 mM) or absence of GP. Bars represent mean values ± standard deviation of triplicate measurements from at least three independent experiments; fold-changes are explicitly indicated. The levels of statistical significance are indicated with ∗∗ *p* < 0.01 (Dunnett’s test). *Concn*., concentration.

**Table 1.** Strains and plasmids used in this work.

| Bacterial strain | Relevant characteristics | Reference |
| --- | --- | --- |
| *Escherichia coli* DH5α λ*pir* | F<sup>-</sup> λ<sup>-</sup> *endA1 glnX44(AS) thiE1 recA1 relA1 spoT1 gyrA96(Nal<sup>R</sup>) rfbC1 deoR nupG Φ80(lacZΔM15) $\Delta$(argF-lac)U169 *hsdR17(rK<sup>-</sup> mK<sup>+</sup>) λpir* lysogen* | Platt et al. [64] |
| *Achromobacter xylosoxidans* SOS3 | Isolated from soil; grows in a medium with GBH as the only source of phosphorus | Masotti et al. [27] |
| *Agrobacterium tumefaciens* CHLDO | Isolated from soil; grows in a medium with GBH as the only source of phosphorus | Masotti et al. [27] |
| *Agrobacterium tumefaciens* GV3101 | Laboratory strain | Holsters et al. [43] |
| Plasmid |  |  |
| PLP1_*P<sub>dummy</sub>* | Reporter plasmid; *oriV*(RSF1010), *P<sub>dummy</sub>*→*mScarlet*; Tet<sup>R</sup> | This work |
| pL1_*P<sub>phnG</sub>* | Reporter plasmid; *oriV*(RSF1010), *P<sub>phnG</sub>*→*mScarlet*; Tet<sup>R</sup> | This work |
| pL1_*P<sub>phnJ</sub>* | Reporter plasmid; *oriV*(RSF1010), *P<sub>phnJ</sub>*→*mScarlet*; Tet<sup>R</sup> | This work |
| pL1_*P<sub>phnC</sub>* | Reporter plasmid; *oriV*(RSF1010), *P<sub>phnC</sub>*→*mScarlet*; Tet<sup>R</sup> | This work |
| pL1_*P<sub>phnD</sub>* | Reporter plasmid; *oriV*(RSF1010), *P<sub>phnD</sub>*→*mScarlet*; Tet<sup>R</sup> | This work |
| pL1_*P<sub>phnE2</sub>* | Reporter plasmid; *oriV*(RSF1010), *P<sub>phnE2</sub>*→*mScarlet*; Tet<sup>R</sup> | This work |
| pL1_*P<sub>duf1045</sub>* | Reporter plasmid; *oriV*(RSF1010), *P<sub>duf1045</sub>*→*mScarlet*; Tet<sup>R</sup> | This work |
| pL1_*P<sub>phnG</sub>ΔB* | Reporter plasmid; derivative of pL1_*P<sub>phnG</sub>* lacking the PhoB binding site (PhoB box) | This work |
| pL1_*P<sub>phnG</sub>ΔF* | Reporter plasmid; derivative of pL1_*P<sub>phnG</sub>* lacking the PhnF binding site (PhnF box) | This work |
| pL1_*FP<sub>phnG</sub>* | Reporter plasmid; derivative of pL1_*P<sub>phnG</sub>* with a copy of *phnF* upstream of the *P<sub>phnG</sub>* promoter | This work |

Tet<sup>R</sup>; tetracycline resistance.

All WCB strains were incubated in MSM medium supplemented with two types of PHTs, i.e., GP or AMPA, using Pi as the phosphorus source in control experiments (**Fig. 2B-E**). The mScarlet fluorescence signal was continuously measured over 16 h (expressed as arbitrary units, A.U.) and normalized to the cell density (estimated as the optical density at 600 nm, OD_600_) to study the transcriptional activity of these constructs. The control strain CHLDO·P*_dummy_* showed no fluorescence in any of the conditions assayed (< 1,550 A.U./OD_600_, **Fig. 2B**). Likewise, strains CHLDO·P*_phnD_*, CHLDO·P*_phnE2_*, and CHLDO·P*_phnJ_* did not yield any detectable mScarlet fluorescence regardless of the presence of GP. This observation indicates that either these three *in*-*silico* identified putative promoters are truncated, or they respond to other signals that may not be captured in a plasmid-based format, where they are isolated from the native genomic context.

Strains CHLDO·P*_phnC_* (**Fig. 2C**), CHLDO·P*_duf1045_* (**Fig. 2D**), and CHLDO·P*_phnG_* (**Fig. 2E**) showed fluorescence levels above the background during growth with either GP or AMPA as phosphorus sources compared to control conditions, where Pi was added to the medium. Strain CHLDO·P*_phnC_* had low fluorescence levels in the presence of Pi (2,270 ± 169 A.U./OD_600_ at 16 h), and no induction was observed when GP was added to the Pi-containing medium (2,542 ± 73 A.U./OD_600_, **Fig. 2C**). In contrast, when this strain was incubated with GP or AMPA as the only phosphorus source, the mScarlet output increased by 76- and 101-fold, respectively. These values were, however, lower than the mScarlet output observed in the absence of any phosphorous additive. A similar behavior was observed for strain CHLDO·P*_duf1045_* (**Fig. 2D**). In this case, the basal fluorescence level in cultures containing Pi was ca. 23-fold higher than that of CHLDO·P*_phnC_* cultivated in the same conditions. The induction of this promoter by GP and AMPA yielded a 4- and 2-fold increase in the mScarlet levels, respectively. The P*_phnG_* promoter, in contrast, responded positively to the presence of PHTs by driving the expression of *mScarlet* to levels above the control without any source of phosphorus. At 16 h, the normalized mScarlet signal increased up to 103,217 ± 3,420 A.U./OD_600_ with AMPA and 149,867 ± 1,567 A.U./OD_600_ with GP (**Fig. 2E**). Taken together, these results suggest that the *phn* cluster spans three PHT-responsive polycistrons: *phnGHIJKLO*, *phnCDE2E1,* and *duf1045 phnMN*, with *phnF* probably forming a monocistronic unit.

When the difference in the maximum mScarlet fluorescence (fold-change) was evaluated among the experiments with strains CHLDO·P*_phnG_*, CHLDO·P*_phnC_*, and CHLDO·P*_duf1045_*, only the P*_phnG_*-biosensor exhibited a significantly increased response to the presence of GP (1.8-fold, **Fig. 2F**). The other two promoters showed a negative response to GP, with a decrease of 30-40% in the fluorescence levels compared to the control experiment, without any added phosphorus source (**Fig. 2F**). These results indicate that the absence of Pi, rather than the presence of a PHT as a source of phosphorus, is a critical factor influencing the expression levels of the P*_phnC_* and P*_duf1045_* promoters. The fact that only the P*_phnG_* promoter responded to the addition of GP to the medium is probably linked to the presence of a PhnF box within its sequence (**Fig. 1C**). Therefore, this promoter was selected as a candidate for constructing a biosensor displaying a positive dose-response behavior to GP.

### Exploring the dose-response profile of a P_phnG_-based WCB

The CHLDO·P*_phnG_* WCB was evaluated in MSM medium supplemented with GP at different concentrations (up to 100 μM) to study the response of the P*_phnG_* promoter to the PHT effector. The fluorescence signal increased linearly over time with the amount of GP when the compound was present at concentrations >2.5 μM (**Fig. 3A**). Above 25 μM GP, the system apparently became saturated and no longer responded to changes in the PHT concentration. This effect was clearly noticeable when the fluorescence fold-changes were calculated at 16 h (**Fig. 3B**). The mScarlet fluorescence was 1.8-fold higher than in the control experiment when the GP concentration was 25, 50, or 100 μM. Significant differences in the fluorescence fold-change were observed at GP concentrations as low as 5 μM, although the increment in the biosensor output was relatively modest (1.2-fold). Although the response of the P*_phnG_* promoter to GP does not follow a strong correlation with the PHT concentration, these results provide a basis for further optimization of the biosensor.

**Figure 3.**
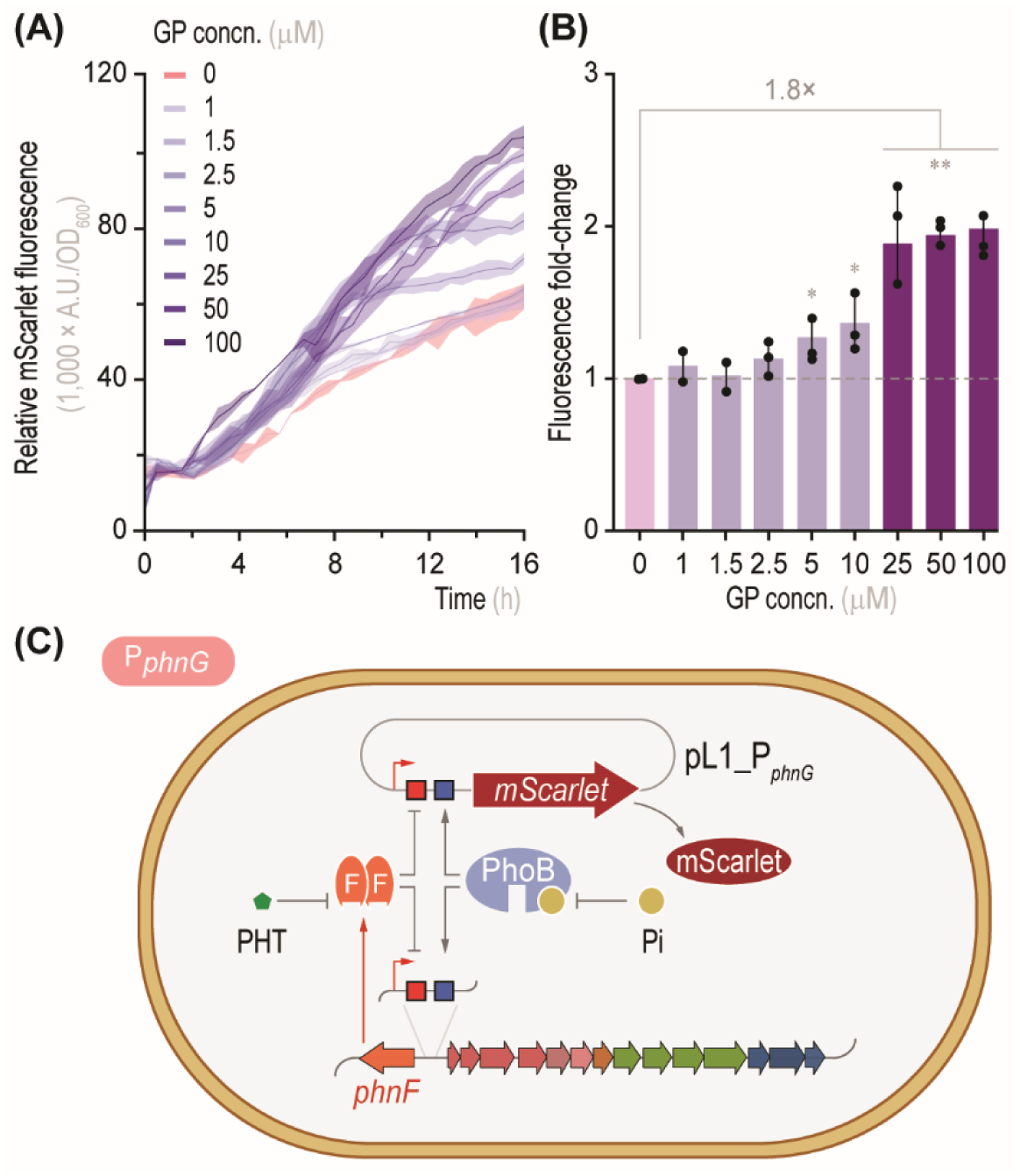
Performance of the P*_phnG_*-based whole-cell biosensor exposed to different GP levels. **(A)** *A*. *tumefaciens* CHLDO, transformed with plasmid pL1_P*_phnG_*, was incubated under the same conditions indicated for **Fig. 2** in the presence of different GP concentrations (*concn*.). Shade lines represent standard deviations for triplicate measurements from at least two independent experiments. **(B)** Whole-cell biosensor response to increasing GP concn. Fold-changes were calculated using the maximum mScarlet output after 16 h of incubation relative to the control experiment without GP; bars represent mean values of the fold-change ± standard deviation from three independent experiments. The levels of statistical significance are indicated with ∗ *p* < 0.05 and ∗∗ *p* < 0.01 (Dunnett’s test). **(C)** Proposed regulatory circuit underlying the P*_phnG_*-based whole-cell biosensor. The PhoB binding box (in blue) becomes active in the absence of phosphate (Pi), whereas the PhnF binding box (in red) is derepressed in the presence of the phosphonate (PHT). This dual-regulation system is expected to yield a red fluorescent signal with no Pi but in the presence of PHT compounds.

The poor performance of this first-generation WCB seems to be connected to the high background fluorescence detected in the absence of GP (ca. 60,000 A.U./OD_600_ after 16 h of incubation). The elevated basal reporter expression limits the sensitivity and dynamic range of the WCB in response to varying PHT concentrations. Based on these observations, we posit that the dual regulation of the P*_phnG_* promoter could be a target for optimizing the biosensor. In addition to the known transcriptional control exerted by the PhnF repressor, the P*_phnG_* promoter is likely subjected to the regulation by the PhoB activator, as previously described for *E. coli* [42] and *Mycobacterium smegmatis* [23]. PhnF and PhoB have been proposed to act as the major players mediating the expression of the *phn* cluster, activating the transcription of the corresponding genes in the absence of Pi or in the presence of a PHT effector, respectively [17]. **Fig. 3C** summarizes the PhnF- and PhoB-dependent regulatory mechanisms expected in strain CHLDO·P*_phnG_*. The results obtained with this first set of WCBs highlight their potential to be employed for GP detection; however, additional engineering is needed to optimize detection capabilities.

### The PhoB and PhnF binding sites upstream of the P_phnG_ promoter are critical for GP detection

To unveil the role of the PhnF and PhoB regulators in the response of the of P*_phnG_* promoter to PHTs, we conducted additional genetic analyses of the promoter region by creating two versions of the pL1_P*_phnG_* plasmid (**Fig. 4**). In one of these constructs, the PhoB box was removed (giving rise to plasmid pL1_P*_phnG__*_Δ*B*_, **Fig. 4A**) to circumvent the Pi-dependent regulation of the promoter. In the other variant, the PhnF box was deleted (plasmid pL1_P*_phnG__*_Δ*F*_, **Fig. 4C**), which is expected to prevent the repression exerted by the PHT-dependent regulator. The corresponding plasmids were transformed into strain CHLDO, and the resulting WCBs were grown as indicated above, adding increasing concentrations of GP in the medium.

**Figure 4.**
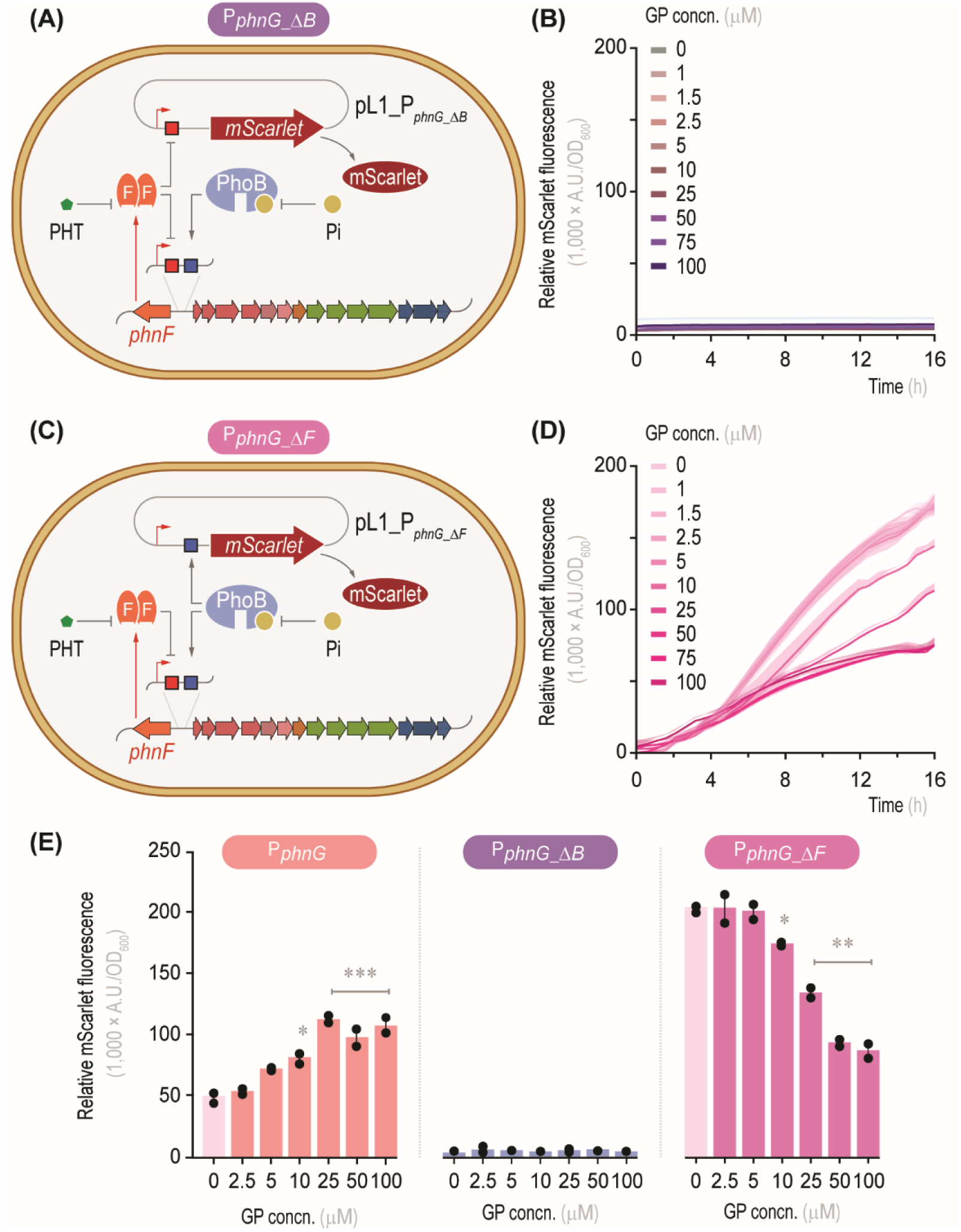
Exploring the dual PhoB and PhnF regulation of the P*_phnG_* promoter. **(A)** Regulatory circuit underpinning the P*_phnG_*__ΔB_-based whole-cell biosensor. Plasmid pL1_P*_phnG_*__ΔB_ carries a deletion in the PhoB box (highlighted in blue), which renders the system responsive to PhnF as the only regulatory signal. **(B)** *A*. *tumefaciens* CHLDO, transformed with plasmid pL1_P*_phnG_*__ΔB_, was incubated under the same conditions indicated for **Fig. 2** in the presence of different GP concentrations (*concn*.). The biosensor output is expressed as arbitrary fluorescence units (*A.U.*) relative to the optical density at 600 nm (OD_600_). Shaded lines represent standard deviations for triplicate measurements from at least three independent experiments. **(C)** Regulatory circuit underpinning the P*_phnG_*__ΔF_-based whole-cell biosensor. Plasmid pL1_P*_phnG_*__ΔF_ carries a deletion in the PhnF box (highlighted in red), which renders the system responsive to PhoB as the only regulatory signal. **(D)** *A*. *tumefaciens* CHLDO, transformed with plasmid pL1_P*_phnG_*__ΔF_, was incubated under the same conditions indicated for **Fig. 2** in the presence of different GP concns. The biosensor output was quantified as explained above. **(E)** Maximum output for the whole-cell biosensors harboring the P*_phnG_*__ΔB_ and P*_phnG_*__ΔF_ constructs after 16 h in the presence (1.5 mM) or absence of GP. The original P*_phnG_* construct was included for comparison. Bars represent mean values ± standard deviation of triplicate measurements from at least three independent experiments. The levels of statistical significance are indicated with ∗ *p* < 0.05, ∗∗ *p* < 0.01, and *** *p* < 0.001 (Dunnett’s test).

The absence of the PhoB box upstream of the P*_phnG_* promoter mediated a complete loss of the mScarlet signal under all tested conditions (**Fig. 4B**). These results indicate a key role for this DNA motif in the recognition by the PhoB activator and underscore the involvement of the Pho regulon as a key factor modulating the expression of the *phn* gene cluster. In contrast, eliminating the PhnF box inverted the responsiveness of the P*_phnG_* promoter to GP. After 16 h of incubation, the fluorescence signal reached by strain CHLDO-P*_phnG__*_Δ*F*_ in the absence of GP was the highest observed (185,879 ± 4,270 A.U./OD_600_, **Fig. 4D**). These values are ca. 3-fold higher than those observed in the strain transformed with the original pL1_P*_phnG_* plasmid grown under the same conditions (64,012 ± 4,033 A.U./OD_600_, **Fig. 3A**). No substantial difference in the mScarlet fluorescence was observed when the CHLDO·P*_phnG__*_Δ*F*_ WCB was incubated in the presence of GP up to 5 μM. These findings suggest that the PhnF box is recognized by the chromosomally-encoded repressor, highlighting the importance of such regulatory layer for the detection and quantification of PHTs. The addition of GP at 10 μM decreased the maximum fluorescence values relative to the control (0 μM GP) by 15%, and increasing the PHT concentration to 25, 50, and 100 μM further reduced the biosensor output by 35%, 55%, and 60%, respectively (**Fig. 4E**). These results suggest that GP degradation in the WCB strain might be releasing Pi (**Fig. 1B**), which in turn partially blocks the activity of the PhoB activator. Since PhoB positively modulates the transcriptional response of the P*_phnG_* promoter, a higher Pi concentration results in a reduced fluorescence output.

To explore the extent of this regulation in related bacterial species, we introduced the relevant sensor plasmids into another *Agrobacterium* strain, *A. tumefaciens* GV3101 (**Table 1**). *A. tumefaciens* GV3101 is a laboratory-adapted strain, widely used as a vector for plant transformation [43]. The corresponding WCB strains, termed GV3101·P*_dummy_* (used as a negative control), GV3101·P*_phnG_*, GV3101·P*_phnG__*_Δ*B*_, and GV3101·P*_phnG__*_Δ*F*_ (**Table S4** in the Supporting Information) were incubated as explained above with varying GP concentrations. Similar to the results observed in experiments with *A. tumefaciens* CHLDO, strains GV3101·P*_dummy_* and GV3101·P*_phnG__*_Δ*B*_ did not exhibit mScarlet fluorescence under any condition (**Fig. S1** in the Supporting Information). Strain GV3101·P*_phnG__*_Δ*F*_ showed maximum fluorescence levels in the absence of GP, with a negative trend as the PHT concentration increased. Strain GV3101·P*_phnG_*, in contrast, displayed a slight increase in the mScarlet output at high (25 and 100 μM) GP concentrations. These results mirror the behavior of the WCBs based on strain CHLDO, indicative of a conserved mechanism for the transcriptional regulation exerted by PhnF and PhoB across *Agrobacterium* species. The fact that strain CHLDO, a highly efficient GP degrader, showed higher fluorescence signals than strain GV3101 in response to GP suggests that specific metabolic and regulatory adaptations to PHT degradation may have been lost in laboratory-domesticated strains.

### Optimization of the WCB towards quantifying PHTs

The lack of response of the biosensor to GP observed when the PhnF box was removed suggested that titrating the amount of regulator available to bind the PhnF motif is a key parameter for optimization. We engineered another version of the P*_phnG_*-based WCB that bears an extra *phnF* copy upstream of the P*_phnG_* promoter (**Fig. 5A**). In the resulting plasmid, termed pL1_FP*_phnG_* (**Table 1**), the *phnF* gene retains its native promoter, and is placed in the opposite direction with respect to the rest of functional elements in the construct. Plasmid pL1_FP*_phnG_* was introduced in strain CHLDO, giving rise to CHLDO·FP*_phnG_*, and the behavior of this new WCB was analyzed in the presence of GP concentrations ranging from 0 to 150 μM. An increase in the mScarlet signal with the GP concentration was observed in these experiments, noticeable even to the naked eye (**Fig. 5B**).

**Figure 5.**
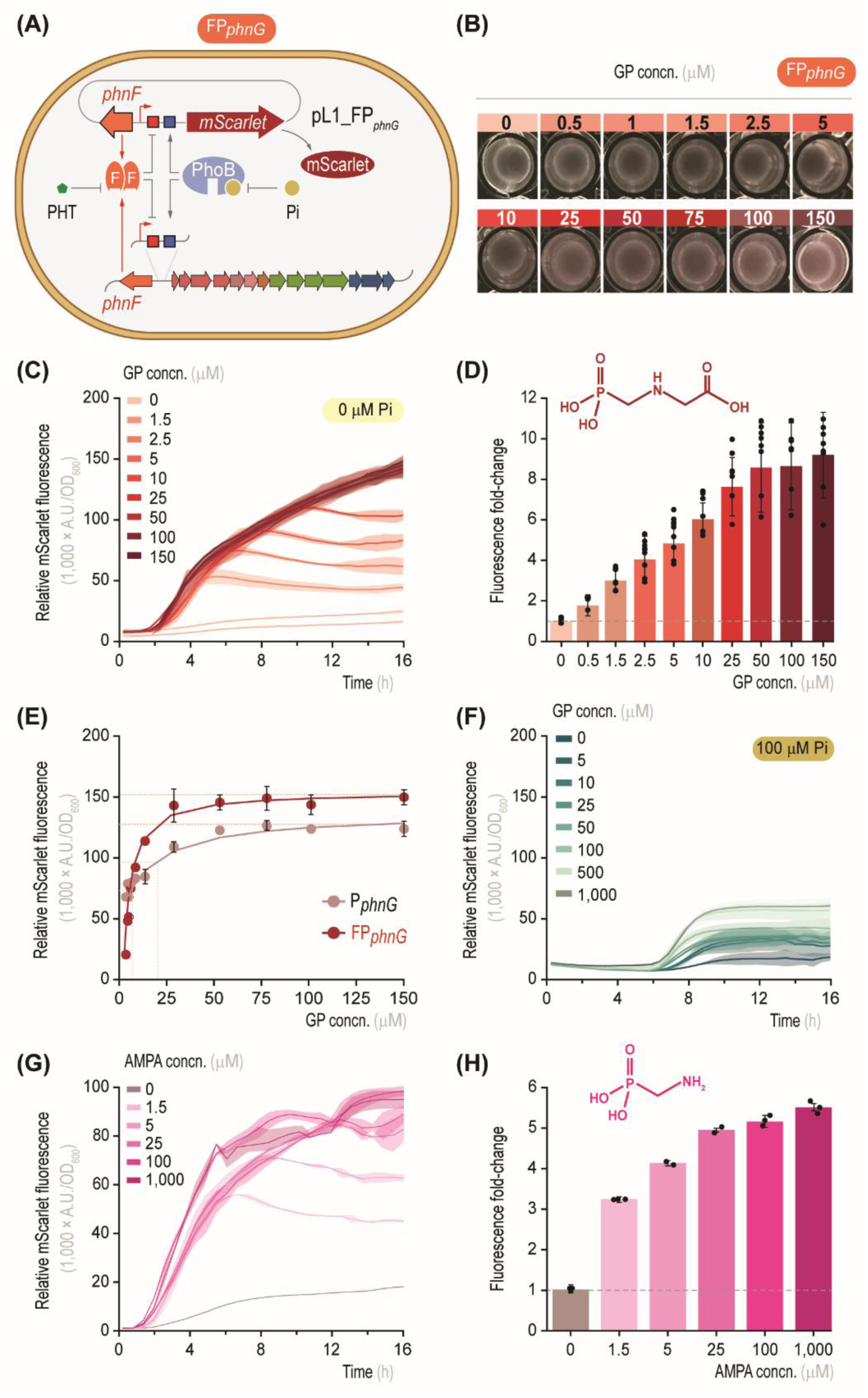
PhnF titration increases the output and sensitivity of the P*_phnG_*-based whole-cell biosensor. **(A)** Engineering PhnF dosage by adding a copy of *phnF* in plasmid pL1_FP*_phnG_*. The orientation with respect to the P*_phnG_* promoter and the transcriptional regulation of the *phnF* gene was kept as in its native context. **(B)** Cultures of *A*. *tumefaciens* CHLDO, transformed with plasmid pL1_FP*_phnG_*, were incubated under the conditions indicated for **Fig. 2** with different GP concentrations (*concn*.). The microtiter well plates were photographed after 24 h. **(C)** *A*. *tumefaciens* CHLDO, transformed with plasmid pL1_FP*_phnG_*, was incubated under the conditions indicated above with different GP concns. and no phosphate (Pi) added. The biosensor output is expressed as arbitrary fluorescence units (*A.U.*) relative to the optical density at 600 nm (OD_600_). Shaded lines represent standard deviations for triplicate measurements from at least three independent experiments. **(D)** Whole-cell biosensor response to increasing GP concns. Fold-changes were calculated using the maximum mScarlet output after 16 h of incubation relative to the control experiment without GP; bars represent mean values of the fold-change ± standard deviation from four independent experiments. In all cases, the differences between the control experiment and those added with GP were statistically significant with ∗ *p* < 0.05 (Dunnett’s test). **(E)** Dose-response functions for the optimized FP*_phnG_*- and the parental P*_phnG_*-based whole-cell biosensor. Datapoints represent mean values of the maximum mScarlet fluorescence ± standard deviation from four independent experiments. **(F)** Performance of the FP*_phnG_*-based whole-cell biosensor in the presence of Pi. Culture conditions and biosensor output were as indicated for panel **(C)**. **(G)** Testing the FP*_phnG_*-based whole-cell biosensor for detection of (aminomethyl)phosphonate (AMPA). Cells were incubated under the same conditions indicated for **Fig. 2** in the presence of different AMPA concns. with no phosphate (Pi) added. The biosensor output is expressed as *A.U.* relative to OD_600_. Shaded lines represent standard deviations for triplicate measurements from at least three independent experiments. **(H)** Whole-cell biosensor response to increasing AMPA concns. Fold-changes were calculated using the maximum mScarlet output after 16 h of incubation relative to the control experiment without AMPA; bars represent mean values of the fold-change ± standard deviation from four independent experiments. In all cases, the differences between the control experiment and those added with GP were statistically significant with ∗ *p* < 0.05 (Dunnett’s test).

The CHLDO·FP*_phnG_* WCB produced virtually no fluorescence in the absence of GP (**Fig. 5C**), as evidenced in the corresponding control culture (**Fig. 5B**). This behavior is different to the results obtained with strain CHLDO·P*_phnG_* (**Fig. 3B**), which had a high mScarlet output in absence of the effector molecule. In contrast, the mScarlet signal emitted by strain CHLDO·FP*_phnG_* increased steadily with the GP concentration (**Fig. 5C**), with a maximum fluorescence value of 150,567 ± 2,106 A.U./OD_600_ after 16 h of incubation. When the relative mScarlet levels were normalized to the control experiment (no GP added), a linear relationship was observed between the fluorescence fold-change and the GP concentration (**Fig. 5D**). Adding the PHT at 0.5 μM almost doubled the fluorescence fold-change, an increase that was achieved with the original construct (plasmid pL1_P*_phnG_*) only when GP was added at 100 μM (**Fig. 3B**). The response of the CHLDO·FP*_phnG_* WCB appeared to saturate at GP concentrations >50 μM, with a fold-change in the mScarlet fluorescence of 8.7 ± 1.2 (**Fig. 5D**). These results indicate that the overall biosensor performance was substantially enhanced by manipulating the levels of PhnF titration.

A typical inducer-dose response was obtained when the maximum fluorescence output by the WCB was plotted against the GP concentration (**Fig. 5E**). The values for the dynamic range (fold-change) obtained through fitting were 8.9 ± 0.5 and 2.1 ± 0.1 for strains CHLDO·FP*_phnG_* and CHLDO·P*_phnG_*, respectively (**Table 2**). The optimized biosensor displayed a 4-fold reduction in the leakiness (i.e., background fluorescence) when compared to the original construct in which no *phnF* copy was included (**Table 2**). The sensitivity of the biosensor to PHT effectors decreased from 17.3 to 4.2 μM GP, with a practical detection range of 0.5 to 50 μM. Hence, addition of an extra *phnF* copy lowered the basal expression of the system due to a tighter transcriptional repression, enabling a lower detection limit than that of the original biosensor.

**Table 2.** Response parameters for the original and the optimized whole-cell biosensor.

| WCB strain | Leakiness | Maximum<br>fluorescence | Dynamic range<br>(fold-change) | Sensitivity<br>(μM) | *n* |
| --- | --- | --- | --- | --- | --- |
| CHLDO-$P_{phnG}$ | 64,012 ± 4,033 | 130,457 ± 4,724 | 2.1 ± 0.1 | 17.3 ± 6.4 | 1.10 ± 0.12 |
| CHLDO-FP$_{phnG}$ | 16,928 ± 1,868 | 150,567 ± 2,106 | 8.9 ± 0.5 | 4.2 ± 0.3 | 1.02 ± 0.03 |

4 The parameters used for adjusting the model are described in Materials and Methods.

These features were likewise verified when the corresponding sensor plasmids were moved to *A. tumefaciens* GV3101 as an alternative host (**Fig. S2** in the Supporting Information) and tested against GP. Strain GV3101·FP*_phnG_* exhibited a higher sensitivity and wider dynamic range than GV3101·P*_phnG_* (**Fig. S2A**), suggesting that the optimized biosensor plasmid can be transferred to other environmental bacteria for detection of PHTs. While the minimum fluorescence levels were similar for both WCBs, the maximum mScarlet signal increased to 57,228 ± 2,583 A.U./OD_600_ in strain GV3101·FP*_phnG_* (**Fig. S2B**). This mScarlet output is lower than that of *A. tumefaciens* CHLDO under the same conditions (**Table 2**), underscoring strain-specific differences in GP processing and transcriptional regulation of the *phn* genes. One such regulatory layer, common to different bacterial GP degraders, is the modulatory effect of Pi. To investigate the impact of Pi on the biosensor response, strain CHLDO·FP*_phnG_* was incubated in MSM supplemented with 100 μM Pi and varying concentrations of GP (**Fig. 5F**). The response time of the system (i.e., the time until the fluorescence signal could be distinguished from the background) increased in all instances, in a range between 6.5-8.7 h compared to 1-2.25 h in the condition without Pi. The addition of Pi also reduced the leakiness of the WCB by ca. 2-fold. As a result, these modifications enabled the detection of GP in the high range of concentrations (up to 1 mM), which was not possible in the condition without Pi (**Fig. 5C**). This behavior was also verified in strain GV3101·FP*_phnG_* (details not shown). Taken together, these results highlight the versatility and portability of the optimized biosensor for GP detection across a range of operational conditions.

We evaluated the CHLDO·FP*_phnG_* WCB for detection of other PHTs. Our previous results showed that CHLDO·P*_phnG_* cells respond to the presence of 1.5 mM AMPA (**Fig. 2E**). Hence, the optimized WCB was incubated in the presence of increasing AMPA concentrations. In these experiments, AMPA could be detected within a 1.25-1,000 μM range (**Fig. 5G**), and at the highest PHT concentration the fluorescence fold-change was 5.6 ± 0.4 (**Fig. 5H**). These results highlight the versatility of the optimized biosensor for the detection of PHTs, as demonstrated with GP and AMPA. The next step was testing the WCB in setups compatible with GP detection in a setup relevant to agricultural applications.

### Applying the CHLDO·FP_phnG_-based WCB for detection of PHTs in laboratory and environmental samples

We explored whether the optimized WCB could be applied in an agriculture-relevant context. Although previous experiments utilized pure PHT effectors, e.g., commercially available GP and AMPA, practical applications involve GP-based herbicides (GBHs). GBHs generally consist of a glyphosate salt combined with other components, including surfactants and preservatives, which are necessary to stabilize the herbicide formulation and facilitate plant penetration. Roundup™, extensively used on genetically-modified corn, soy, and cotton crops resistant to the herbicide, is a common GBH initially developed by Monsanto in the 1970s. Hence, we evaluated whether the CHLDO·FP*_phnG_*-based WCB could detect GP in Roundup™ ControlMax, containing 79.2% (w/w) GP as the monoammonium salt. The biosensor was evaluated using various GBH concentrations (ranging from 0 to 30 μM GP, reflecting levels typically encountered in some field applications). The mScarlet outputs were comparable to those obtained with pure GP (**Fig. S3A** in the Supporting Information). The GP concentrations in these samples were also quantified using high-performance liquid chromatography (HPLC, **Fig. S3B**). An excellent correlation was observed between the GP concentrations determined by the biosensor and those measured using analytical methods. These results indicate that the biosensor strain can be used for PHT detection even in the presence of other components usually found in GBH formulations.

Building on these results, we evaluated the use of the WCB for detecting GP under three scenarios (**Fig. 6**), including microbial PHT degradation, monitoring GBH levels in contaminated soils, and *in situ* testing of environmental samples. In the first case, we characterized GP degradation by *Achromobacter xylosoxidans* SOS3, a Gram-negative species isolated from GBH-contaminated soils previously described to grow on GP as the main source of phosphorous [27]. *A. xylosoxidans* SOS3 was grown in MSM added with 1.5 mM GBH, and samples from these cultures were taken at 72- and 96-h of growth. After spinning down the biomass, the residual GP concentration was quantified with both the CHLDO·FP*_phnG_* WCB and HPLC (**Fig. 6A**). No substantial GP degradation was observed over the first 72 h of incubation, with the PHT concentration reaching 1,232 ± 187 μM, still within the range of the initial levels (1,503 ± 318 μM). The GP concentration substantially decreased after 96 h, reaching 376 ± 36 μM (**Fig. 6B**), with a close correlation between the WCB- and the HPLC-based determinations. These results suggest that bacterial PHT degradation is highly dependent on the amount of biomass, since GP is slowly consumed if the cell density remains below a threshold, as previously reported [27]. These observations indicate that the WCB could be used for quantifying herbicides in laboratory setups as an efficient alternative for traditional methods for GBH detection, eliminating the need for the derivatization step required in HPLC analysis.

**Figure 6.**
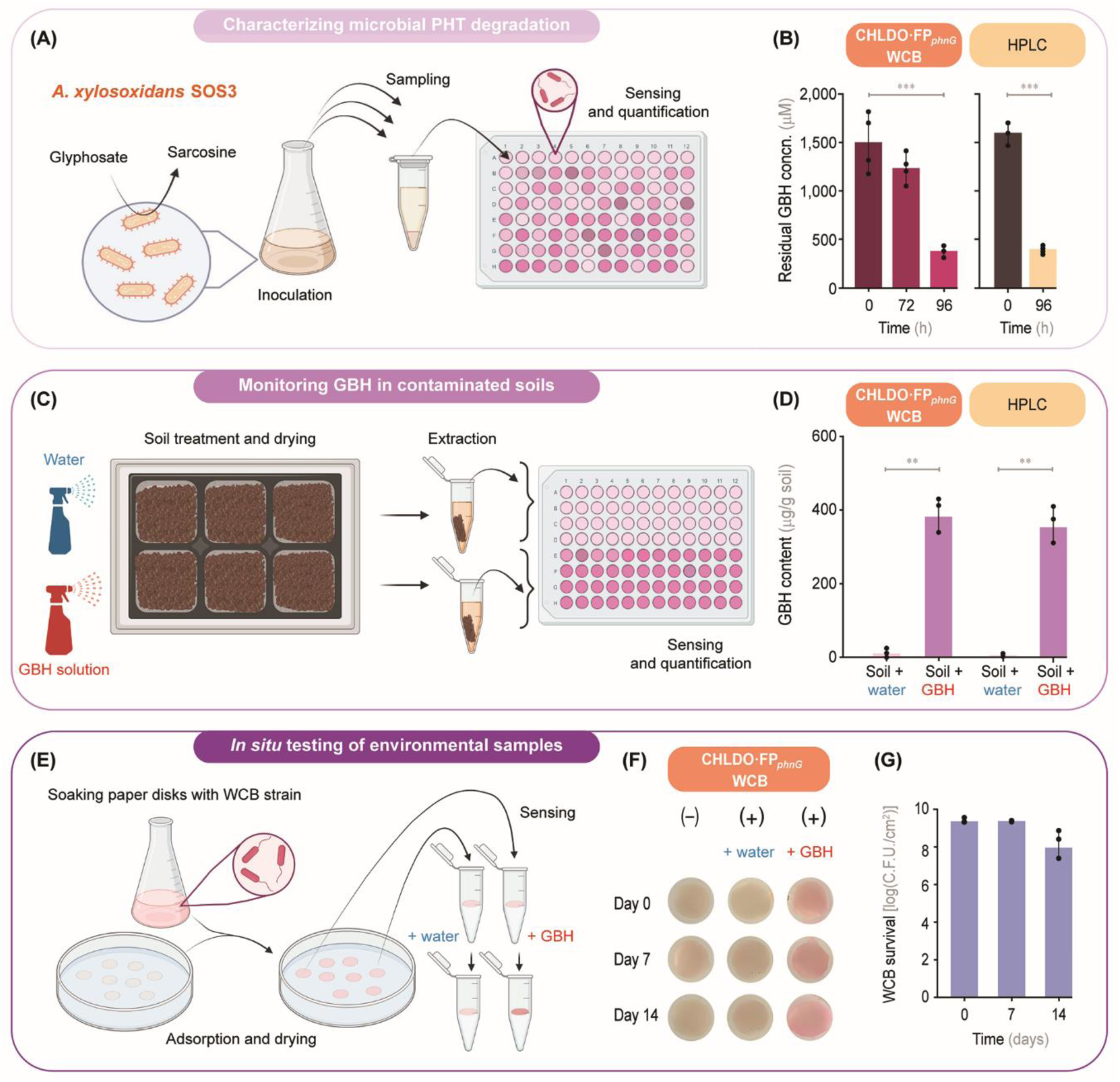
Applications of the FP*_phnG_*-based whole-cell biosensor. **(A)** Exploring microbial phosphonates (PHT) degradation. *Achromobacter xylosoxidans* SOS3 can degrade GBH (a glyphosate-based herbicide) in liquid culture, releasing sarcosine as the main end-product. *A*. *xylosoxidans* SOS3 was grown at 30°C in MSM medium supplemented with 1.5 mM GBH; samples were taken at the specified times throughout a 96-h incubation period. **(B)** The residual GBH concentration (concn.) in these aliquots was quantified using the FP*_phnG_*-based whole-cell biosensor (WCB); HPLC-based quantification was done in parallel in the same samples. **(C)** Quantification of GBH in contaminated soils. Soil samples, disposed in pots, were sprayed with a GBH solution (or water, as a control) and dried for 3 days at 37°C. Soil samples (50 mg) were suspended in 500 μL of water and, after a 30-min incubation with shaking, the tubes were centrifuged, and the supernatants were collected for GP detection with the FP*_phnG_*-based WCB. **(D)** GBH quantification in soil samples sprayed with a 4 mM GBH solution or water with both the WCB and by HPLC. In panels **(B)** and **(D)**, the bars represent mean values ± standard deviation of triplicate measurements from at least three independent experiments. The levels of statistical significance are indicated with ∗∗ *p* < 0.01 and *** *p* < 0.001 (Dunnett’s test). **(E)** Detecting GBH in agar pads with the FP*_phnG_*-based WCB. Filter discs were soaked with a saturated culture of *A*. *tumefaciens* CHLDO carrying plasmid pL1_FP*_phnG_*, dried, and kept for up to 14 days at 4°C. The filter discs were placed onto MSM agar and added with a 20-μL drop of 1.5 mM GBH (or water, as control); the samples were photographed after a 48-h incubation at 30°C. **(F)** Representative pictures of the WCB-embedded filter disks exposed to water or the GBH solution after 0, 7, and 14 days. Filter discs that have not been soaked with the biosensor strain, indicated as (–), are included as a control. **(G)** Bacterial survival in the filter-disk format. WCB cells were recovered at the same data points used in the GBH detection assay and plated onto a rich medium for colony forming units (C.F.U.) counting after 24 h. The bars represent the average value of three independent replicates ± standard deviations.

In the second application, we explored the ability of the CHLDO·FP*_phnG_* WCB to detect GBHs in environmental samples (**Fig. 6C**). To this end, soil samples were treated with a solution of GBH at recommended field application doses (ca. 300-500 μg/g soil). As a negative control, soil samples were sprayed with water. After aqueous extraction, the supernatant samples were incubated with the biosensor and assessed by HPLC as explained above. As expected, no GP was detected in the negative control. Soil samples treated with GBH, however, had a GP level of 390 ± 18 μg/g soil, with results consistent with those obtained by HPLC (**Fig. 6D**). Residual GP has been found in soils treated with Roundup^TM^ in a range typically spanning between 2 and 800 μg/g soil [44]. Thus, the WCB developed here could be applied for detection of GP in environmental samples.

Considering the importance of on-site detection and the need for cost-effective devices to assess the presence of GBH pollutants, we explored *in situ* analysis of environmental samples with our biosensor strain. In the third application example, the CHLDO·FP*_phnG_* WCB was inoculated on paper disks, and after drying, the disks were placed onto MSM and treated with a drop of 1.5 mM GP (**Fig. 6E**). A drop of water was used in negative control experiments. Visual inspection of the disks after 48 h of incubation revealed a clear difference in color between the control experiment and the filter disks exposed to GBH (**Fig. 6F**). These results support the use of the WCB in a low-technology setup for the detection of GBHs in environmental samples when fluorescence detection is not readily available. We also studied the stability of WCB cells upon prolonged storage. In these experiments, the biomass bound to the filter disks was recovered after 7 and 14 days of storage at 4°C. The corresponding bacterial suspension was plated, and the survival of the WCB strain was estimated by counting colonies (colony forming units, C.F.U.) and comparing the counts from a freshly prepared filter disk. A consistent number of active bacteria was recovered at the three time points assayed (>10^8^ C.F.U./cm^2^, **Fig. 6G**), with a slight, non-significant decrease in the bacterial count observed at 14 days. Hence, the results in this section expose the use of the WCB for detection of PHTs across different operating conditions that include field-relevant agricultural applications.

## Conclusion

GP, the most widely used non-selective, broad-spectrum herbicide globally, has well-documented adverse effects on the environment and human health. Strategies to detect and control the presence of GBHs and related compounds in the environment have become increasingly important as the use of GBHs continues to rise. On this background, we developed an affordable, simple, and accurate WCB platform by harnessing the native regulatory circuit of *A*. *tumefaciens* CHLDO, a GP-degrading bacterium isolated from contaminated soils. To our knowledge, this is the first instance where a TF is used to sense PHTs. The analysis of operator sequences within the *phn* gene cluster of *A*. *tumefaciens* CHLDO enabled the isolation of several candidate GP-responsive promoters. Among these, the P*_phnG_* promoter was selected to drive the transcription of a fluorescent protein (mScarlet) gene for PHT sensing. Incorporating this construct into *A*. *tumefaciens* CHLDO resulted in a first-generation WCB, which underwent several rounds of optimization. Adding an additional copy of *phnF* to the construct enhanced the repression of the promoter, minimizing its leakiness and significantly improving the dynamic range. This modification enhanced the detection capability for measuring GP levels, enabling applications in both laboratory and environmental setups. Additionally, FP*_phnG_* lacks specificity for GP and can also detect a primary GP-degradation product, AMPA.

Further optimization could enhance the sensor selectivity for GP and other PHT effectors, e.g., through promoter (P*_phnG_*) and protein (PhnF) engineering. These strategies have been previously applied to other biosensors [32]. For example, a WCB strain was optimized for ligand specificity for mercury (Hg) detection. GolS, a transcription factor of *Salmonella enterica* serovar Typhimurium that activates the response to Au^+^, was subjected to protein engineering to add a third ligand for metal (Hg^2+^) coordination. To further improve specificity towards Hg^2+^, the metal-binding loop of GolS was replaced with a variant from *Pseudomonas aeruginosa* Tn-501. The final version of the engineered GolS-based WCB could detect Hg^2+^ in the nM range, even in the presence of other heavy metals [45]. The crystal structure of the PhnF TF from *M*. *smegmatis* revealed that this transcription factor is a homodimer in its active form, with a helix-turn-helix N-terminal domain and a C-terminal domain that can bind to a yet unidentified ligand [23]. Mutational analysis of key amino acids in the C-terminal domain of PhnF may improve both specificity and sensitivity towards PHTs.

Besides changes in protein sequence that modify the TF structure, promoter engineering to modulate the expression of TF or reporter genes has also been widely utilized for biosensor optimization. For example, optimizing a WCB for Au^3+^ detection involved mutating sequences at the 3’-end and three operator positions of the P*_cupA_* promoter, which responds to the CupR TF [46]. Another relevant case is a WCB for arsenic, which depends on the ArsR TF. In this case, the P*_ars_* promoter was redesigned to achieve a tightly regulated expression, with maximum repression and optimal fold-change, by maximizing the overlap between the promoter core region and the ArsR binding site [47]. A biosensor based on the PHT-binding protein PhnD, part of the PHT transport system from *E*. *coli*, has been employed for PHTs detection. In this biosensor, the PhnD protein has been engineered so that conformational changes upon substrate binding alter the fluorescence emitted by an acrylodan fluorophore attached to the protein near the hinge. However, this PhnD-based biosensor was not reactive to GP and was further engineered to increase its affinity for the PHT effector. Mutations in the glutamic acid 177 to asparagine and the deletion of the last 6 amino acids resulted in a change in the GP affinity, enhancing the biosensor sensibility by >150-fold. However, this new version lacks specificity, as it can bind AMPA, Pi (PO_4_^3–^), and arsenate (AsO_4_^3–^) with similar affinity than that of GP [48].

An analysis of sensing methods for PHTs reported in recent literature (**Table S5** in the Supporting Information) indicates that the TF-based WCBs constructed here compare favorably with other chemical- and fluorescence-detection methods. Their properties, in terms of both analytical performance and key features (e.g., limit of detection, dynamic ranges, and equipment needed for sensing), set a benchmark for *in vivo* detection of PHTs. The results reported here could also aid the development of whole-cell and cell-free biosensors for other applications, including automated detection of contaminants [49] and growth-coupled strategies for assimilation of GP and other PHTs [50]. Meanwhile, the *Agrobacterium* WCBs in this study are useful to monitor environmental samples using cost-effective photometers or even visualized by the naked eye, in contrast to the traditional reliance on spectroscopy and chromatography methods.

## Materials and Methods

### Reagents and culture media

The GBH used in this work was Roundup^TM^ ControlMax, containing 79.2% (w/w) GP in the monoammonium salt form (Monsanto, United States). All concentrations indicated in the experiments using GBH concentration refer to the GP content in the herbicide. GP and AMPA (purity > 98%; Sigma-Aldrich Co., St. Louis, MO, USA) were used for testing and calibration of WCBs. All other chemicals used in this study were of analytical grade. For routine cloning procedures and propagation, bacteria were cultivated in lysogeny broth (LB), containing 10 g/L tryptone, 5 g/L yeast extract, and 10 g/L NaCl [51]. Biosensor experiments were carried out in a mineral salt medium (MSM) containing (in g/L): L-glutamic acid (potassium salt), 12; NH_4_Cl, 2.0; MgSO_4_.7H_2_O, 0.2; and K_2_SO_4_, 0.5. MSM was added with a trace element solution [52], with the following final trace concentrations (in mg/L): FeSO_4_·7H_2_O, 2.5; CaCl_2_·6H_2_O, 10.0; CuSO_4_·5H_2_O, 2.0; H_3_BO_3_,0.06; ZnSO_4_·7H_2_O, 20.0; MnSO_4_·H_2_O, 1.0; NiCl_2_·6H_2_O, 0.05; and Na_2_MoO_4_·2H_2_O, 0.3. MSM medium was supplemented with KH_2_PO_4_ (referred to as Pi in the text), GBH, GP, or AMPA as indicated in the corresponding experiment description.

### Bacterial strains and growth conditions

All bacterial strains used in this study are listed in **Table 1** and **Table S4** in the Supporting Information. Cultures of *A*. *xylosoxidans* SOS3, *A*. *tumefaciens* CHLDO, *A*. *tumefaciens* GV3101, and their engineered derivatives were incubated at 30°C. *E*. *coli* cultures were incubated at 37°C. For routine cloning procedures and overnight cultures to be used as inocula, cells were grown in LB. All liquid cultures were agitated at 200 rpm; solid culture media also contained 15 g/L agar. Whenever needed, chloramphenicol or tetracycline were added at 30 μg/mL or 10 μg/mL, respectively, from filter-sterilized 1,000× stock solutions.

For phenotypic characterization of biosensor strains in microtiter plates, *A*. *tumefaciens* cells were pre-grown in LB. Experiments were performed in MSM supplemented with the different phosphorus sources corresponding to each experimental condition. Pre-cultures were harvested by centrifugation at 4,000×*g* for 8 min, washed with MSM without any phosphorus source, and resuspended in the final medium supplemented tetracycline and GP, GBH, AMPA, or Pi at different concentrations at an OD_600_ of 0.3. Quantification of red (mScarlet) fluorescence was performed in a 96-well plate format in an automated Synergy 2 plate reader (BioTek Instruments; Winooski, VT, USA) [53–55]; the excitation and emission wavelengths (λ*_excitation_*/λ*_emission_*) used were 530 ± 20 nm and 620 ± 20 nm, respectively. The mScarlet measurements were normalized to the OD_600_, registered in the same timelapse every 30 min.

### General cloning procedures and plasmid construction

The DNA constructs described in this work were cloned using Golden Gate assembly [56] unless otherwise stated. The fragments used for cloning were amplified by PCR with Phusion Hot Start II High-Fidelity DNA Polymerase enzyme (Thermo Fisher Scientific Inc). In these cases, 10 ng of DNA and specific oligonucleotides containing at least 25 bp of homology with the destination vector (**Table S3** in the Supporting Information) were used for assembly. After plasmid (backbone) amplification, treatment with the enzyme DpnI (New England BioLabs, Ipswich, MA, USA) was performed for 30 min at 37°C. Next, fragments were purified with a column or from a gel, and a reaction mixture was set with the Gibson Assembly Master Mix by following the supplier’s instructions (New England BioLabs). The reaction was carried out in a thermocycler at 50°C for 15 min. Assembly reactions were dialyzed with MF-Millipore cellulose ester membranes (Merck) and transformed into electrocompetent *E*. *coli* DH5α λ*pir* cells. For colony PCR, the commercial OneTaq™ master mix (New England BioLabs; Ipswich, MA, USA) was used according to the manufacturer’s instructions. The identity and correctness of all plasmids and DNA constructs were confirmed by sequencing.

### Golden Gate Assembly of biosensor plasmids

Golden Gate assembly reactions, in a final volume of 10 μL, were set as follows: 25 fmol of plasmid DNA bearing the antibiotic cassette, 100 fmol of plasmid DNA from other parts, 1 μL T4 DNA ligase buffer, and 1 μL Master Mix Golden Gate BsmbI or BsaI (New England Biolabs) for construction of Level 0 (L0) or Level 1 (L1) plasmids, respectively. The reactions were incubated in a thermocycler with 30 cycles of 5 min at 37°C and 10 min at 16°C, followed by a final incubation at 37°C for 60 min. Finally, an enzyme inactivation step of 10 min at 80°C was performed. Plasmids and oligonucleotides used for introducing *phn* promoters into L0 plasmids are listed in **Table S2** and **S3** in the Supporting Information, respectively. The parts chosen for constructing L1 plasmids belong to Marburg Collection [41], and are indicated in **Table S1** in the Supporting Information.

### Electroporation of *E. coli* and *A. tumefaciens* cells

Electrocompetent cells of *E. coli* and *A. tumefaciens* were prepared as detailed by Warren et al. [57]. Next, 50-μL aliquots of the electrocompetent cell suspensions were thawed and incubated with 50-200 ng plasmid (or dialyzed assembly reactions), kept on ice for 15 min, and then added to 0.2-cm electroporation cuvettes (Bio-Rad) [58]. These reactions were pulsed with a 25 μF, 200 Ω, 2.5 kV, using a Bio-Rad Gene Pulser^TM^ electroporator. Then, *E. coli* or *A. tumefaciens* cells were recovered with 1 mL of LB at 37°C or 30°C for 2 or 4 h, respectively. After this incubation, cells were plated into LB agar supplemented with the corresponding antibiotics. The plates were incubated at 37°C or 30°C until the appearance of colonies.

### *In silico* analysis of *phn* cluster promoter sequences

Analyses of *phn* cluster promoter sequences were performed using BPROM (Softberry Inc., Mount Kisco, NY, USA; http://linux1.softberry.com) [59] to identify the bacterial –10 and –35 sequences and the possible associated TFs. The PhoB binding sites sequences were searched by homology with the motifs described by Yuan et al. [60]. RegPrecise (http://regprecise.lbl.gov) [40] was used for identifying the PhnF binding sequences.

### Fluorescence biosensor analysis

Growth and fluorescence data measured over time was analyzed with QurvE software [61]. Fluorescence data was normalized to growth; data fitting and biosensor dose– response determination was performed using the linear regression model, adapted from Meyer et al. [62], according to Equation (1):

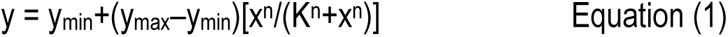

where y is the promoter activity in A.U./OD_600_, *x* is the concentration of the analyte, y_min_ is the leakiness, y_max_ is the maximum fluorescence signal, K is the sensitivity, and *n* is the cooperativity index. This analysis was used to compute leakiness, sensitivity, dynamic range and maximum response; the QurvE software [61] was used for plotting and analyzing fluorescence over time to derive these parameters.

### Analytics

Standard stock solutions of 10 mM GP were prepared in miliQ water. Working standard solutions were prepared at 1 μM by serial dilutions in water, to run 0 to 10 picomoles standard curves (corresponding to 0-2 μM under the conditions assayed). Samples were diluted to fit in this range and derivatized with 9-fluorenylmethoxycarbonyl chloride as described by Wang et al. [63] with some modifications. Briefly, 500 μL of each sample or standard was incubated with 100 μL of water, 100 μL of 0.2 M borate buffer solution (pH = 9), and 100 μL of 1 mg/mL 9-fluorenylmethoxycarbonyl chloride freshly prepared in acetonitrile. After vortexing and incubating the samples for 30 min in the dark, 800 μL of dicloromethane were added, the tubes were vortexed to extract free 9-fluorenylmethoxycarbonyl chloride, and centrifuged for 5 min at 10,000×g. Next, 400 μL of the supernatant were transferred to another tube, 100 μL of methanol were added and after vortexing, samples were filtered by passage through a 0.22-μm syringe nylon filter. Finally, filtered samples were placed in glass HPLC vials. HPLC runs were performed in a Shimadzu LC modular chromatographic system (LC-2050C 3D Plus), equipped with an autosampler (LC-2050) and a FLD (RF-20A). The Shimadzu LabSolutions LC software was used for recording the chromatograms and the analysis of peak areas. The separation of FMOC-derivatives was done on a Shimadzu Shim-pack GIST C18 column (100×3 mm, 3 μm particle size). The FLD was set at 267 nm and 317 nm (excitation and emission, respectively). The mobile phase was composed of 5 mM ammonium acetate (pH = 9, adjusted with NH_4_OH) and methanol. In all cases, 5 μL of each sample were subjected to a 20 min run with the following changes in methanol percentages (v/v): 0 min, 20%; 1.5 min, 20%; 3 min, 70%; 15 min, 70%; 19 min, 20%; and 20 min, 20%; at 0.35 mL/min.

### GP degradation by *A. xylosoxidans* SOS3 measured with a WCB

*A. xylosoxidans* SOS3 was grown in MSM supplemented with GBH as the only phosphorous source. Supernatants of bacterial growth were obtained by centrifugation at specific time intervals throughout a 96-h incubation period at 30°C with shaking at 200 rpm. Next, 3.5 μL of the collected culture supernatant samples were diluted with 1 mL of MSM medium, and 200 μL of this solution were then subjected to incubation with the WCB strains.

### Biosensor detection of GBH-contaminated soil samples

Plastic pots (h=10 cm, d=6 cm) containing 50 g of commercial peat-based potting material were sprayed with 3 mL of a 4 mM GBH solution. As control, other pots were sprayed with 3 mL of water. The pots were incubated for three days at 37°C to allow for soil drying. Then, 50 mg of the topsoil was recovered and suspended in 0.5 mL of miliQ water to dissolve the GP adsorbed to soil particles. After shaking and incubation, the samples were centrifuged twice at 12,000×g for 10 min. Finally, 7.5 μL of these supernatants were diluted to 1 mL of MSM and 200 μL of this solution were incubated with the WCBs as indicated in the text.

### Bacterial immobilization to blotting paper

The WCB strains were grown in LB supplemented with tetracycline with shaking at 200 rpm at 30°C during 24 h. Then, 10 mL of the cell suspension was washed with MSM without any phosphorus source and incubated during 2 h for adaptation. Next, round-shaped blotting sterile papers (ca. 6-mm diameter) were soaked with 50 μL of culture (at an OD_600_ = 1) during 30 min at room temperature for WCB immobilization. The filter disks were dried in sterile conditions and kept at 4°C for up to 14 days. To determine if the immobilized WCB could sense GBH, bacteria-soaked filter disks (0, 7, and 14 days) were placed onto MSM agar and 5 μL of 1.5 mM GBH was added on top of the disks. The coloration was examined after the incubation at 30°C for 48 h. As a negative control, WCB-immobilized paper disks were incubated with 5 μL of sterile miliQ water. Furthermore, to test biosensor survival, colony-forming units (C.F.U.) recovered from immobilized-bacteria paper disks were determined at 0, 7, and 14 days of storage by serial dilution and incubation in LB agar at 30°C for 48 h.

### Statistical analysis

Data analysis was performed using R statistical packages. The reported values are averages of replicates with their respective standard deviations as specified in the figure legends. When relevant, the level of significance of the differences was evaluated using ANOVA of one or two factors, followed by Dunnett’s test.

## Suporting Information

**Table S1.** Gene fragments and synthetic modules used in this work

**Table S2.** Vectors used for Golden Gate assembly of biosensor plasmids

**Table S3.** Oligonucleotides used in this work.

**Table S4.** Whole-cell biosensor strains constructed in this work.

**Table S5.** Summary of recent sensing technologies for GP detection.

**Figure S1 ·** Performance of *A*. *tumefaciens* GV3101-based whole-cell biosensors upon incubation with increasing GP concentrations.

**Figure S2 ·** Performance of optimized *A*. *tumefaciens* GV3101-based whole-cell biosensors.

**Figure S3 ·** Response of the *A*. *tumefaciens* CHLDO·FP*_phnG_* whole-cell biosensor to GBH.

## Acknowledgements

We thank Nicolás Gurdo (DTU Biosustain, Denmark) and Paula Ruffatto (IBR-CONICET, Argentina) for their assistance with HPLC measurements. N.G. and J.O. are staff members and F.M. and N.L. are fellows of Consejo Nacional de Investigaciones Científicas y Técnicas (CONICET). This work was supported by grants from ANPCYT (PICT2020-0838) and MINCYT (B19-CONVE-2021-78800688-APN-SSCI#MCT) to J.O. The financial support from The Novo Nordisk Foundation (grants NNF10CC1016517, NNF18CC0033664, and NNF23OC0083631) to P.I.N. is also gratefully acknowledged.

## Abbreviations

*PHT*: phosphonate
*GP*: glyphosate
*WCB*: whole-cell biosensor
*AMPA*: (aminomethyl)phosphonate
*GBH*: GP-based herbicides
*TF*: transcription factor

