## Supporting Information for "Disentangling the regulatory response of *Agrobacterium tumefaciens* CHLDO to glyphosate for engineering whole-cell phosphonate biosensors"

by

Fiorella Masotti[1‡], Nicolas Krink[2‡], Nicolas Lencina[1], Natalia Gottig[3], Jorgelina Ottado[1*], and Pablo I. Nikel[2*]

[1] Instituto de Biología Molecular y Celular de Rosario, Consejo Nacional de Investigaciones Científicas y Técnicas (IBR-CONICET) and Facultad de Ciencias Bioquímicas y Farmacéuticas, Universidad Nacional de Rosario, Rosario, Santa Fe, Argentina
[2] The Novo Nordisk Foundation Center for Biosustainability, Technical University of Denmark, Kgs. Lyngby, Denmark
[3] Instituto de Procesos Biotecnológicos y Químicos Rosario (IPROBYQ-CONICET-UNR), Rosario, Santa Fe, Argentina

‡ These authors contributed equally.

---

* For correspondence:

**Pablo I. Nikel**
DTU Biosustain, 2800 Kgs. Lyngby, Denmark

**Jorgelina Ottado**
IBR-CONICET, S2000EZP Rosario, Santa Fe, Argentina

---

**Table S1.** Gene fragments and synthetic modules used in this work.

| Description | Sequence (5'→3') |
|---|---|
| 5' Connector<br>**5C5CSR**<br>Golden Gate overhangs | AACA**CGTCTCGAGCG**GGAG |
| **Promoter P$_{dummy}$**<br>Golden Gate overhangs | GGAG**CCCCTGGCGCCCCTT**TACT |
| **Promoter P$_{phnG}$**<br>Golden Gate overhangs<br>**PhoB binding site**<br>**PhnF binding site** | GGAG**GTCACACGCTTGTCATTCATCGACGTTAATTGTAGCTTATATTGTATA**<br>**GTTGTCTATATCAATAGACATTAGAGGTCAAGCG**TACT |
| **Promoter P$_{phnG_\Delta F}$**<br>Golden Gate overhangs<br>**PhoB binding site** | GGAG**GTCACACGCTTGTCATTCATCGACGTTAATTGTAGCTTATATTGTATA**<br>**GTTTAGAGGTCAAGCG**TACT |
| **Promoter P$_{phnG_\Delta B}$**<br>Golden Gate overhangs<br>**PhoB binding site**<br>**PhnF binding site** | GGAG**TCATCGACGTTAATTGTAGCTTATATTGTATAGTTGTCTATATCAATA**<br>**GACATTAGAGGTCAAGCG**TACT |
| **PhnF-Promoter P$_{phnG}$**<br>**PhoB binding site**<br>**PhnF binding site**<br>Golden Gate overhangs | GGAG**TCAATTTTCCACCGTGAATTCGACCGTATCGGCCGAAAAACGGCTGAT**<br>**GGCGTATTGCACCGGCACGCCATCCATATCGGCGTTCAGCGCCTGGGTGACG**<br>**AGCAGGATCGCGCCCGGCGACAATTCGAGATCGGCCAGATCCTGCGTATCCG**<br>**CATGGCTGGCGGAGATGACGGTGGAAATGCGCACGTAATCCGCCACGCCGAG**<br>**CATCTGGAAAGCGGCCGTGATCGAGCCGCTTTTCTGATAGGCCTCGCCTATT**<br>**CCGGCAAAACGGTCGGCCGGGAACCATGTGGTCGAGCGGGATACAGGGCGGT**<br>**TATCCGCCTTGCGCAGCGTTTCGAGGCGCACGAGAGGCGCGCCTGATGGCAC**<br>**TTGCAGTCGCGCCGCCAGATCGGCATTGGCCGGCTCGGTGGTATGGGACAGA**<br>**AGCAGCGCCTCCATTTCCTTCACCTGATCGCCAATGCCCTCCGTAAATCGCG**<br>**TGCGCCGTGAAATCGGAAAACTCAGGCGCTCCTTGCGGGCGATCATCGTGCC**<br>**ACGCCCCTGCATTGGCTCGAGAATGCCTTCACTGGCCAATGCGGCAATGGCG**<br>**CTTCTCACAGTATGGCGGTTGACGCCGAATTTTTCAGCCAGCGCCATTTCCG**<br>**GCGGCAACATGCCGGTTTCGTCGTGATCGCCTGCCGCGATCTCGCCTCTGAT**<br>**CCGGTCGGCGATCTGCCGCCAGAGCGCCACGCCGTTCTTTCTTTTCATCGCT**<br>**GCCGCAGGGCTCATGTCACACGCTTGTCATTCATCGACGTTAATTGTAGCTT**<br>**ATATTGTATAGTTGTCTATATCAATAGACATTAGAGGTCAAGCG**TACT |
| **Promoter P$_{phnC}$**<br>Golden Gate overhangs | GGAG**AATCTCACCTCTGGCAAGGCAAGTGCAAATCCCGGGGCGAAACCCCGG**<br>**GATTTTTGTCGATACACTCTAAAATGCCGGCGAATTTCAACGCCCTGACATG**<br>**CCTCTCGAGAGCTTCGTCGGCGAACAGGTGGACGCCAAGCCCGGGGAAATCT**<br>**GTAATATTTTATTCATCAAACTGACATTCCCACTTCACGCGCCGCTGTTACG**<br>**TCGAACCCCGTCAAAAGAAGATTGGACGGGGAA**TACT |
| **Promoter P$_{phnD}$**<br>Golden Gate overhangs | GGAG**GGAACCGCTCAGCATCGTTTGCGCCCGCGTCAAGGGCCGGTATTTCAA**<br>**AACAAACTCCGGCTCGCCGGGAAACAGGAGAAAGTCC**TACT |
| **Promoter P$_{phnE2}$**<br>Golden Gate overhangs | GGAG**TCCTGACCCACCGTTCCGAAAGCGGCGGTGCCTTGCGGCATCGCCGTT**<br>**TTTGCCAATAAAGGAAAGCGGC**TACT |
| **Promoter P$_{phnJ}$**<br>Golden Gate overhangs | GGAG**CTCAGCGGCTGCCCTGCTCACCAAACCAAAGGGCGCAAAAGACGGTCG**<br>**ATGCTGCAAAGACAGCAAAGAACACGGCTGCAAGAATTCCAATCTCCATGAG**<br>**CTTGTACTCCTTTTTTCACAAGGCTAACGCGCA**TACT |

| | |
|---|---|
| **Promoter $P_{duf1045}$**<br>Golden Gate overhangs | GGAGGCTTTGATGGGGTGGGATGACGAAAGTCGCAACGACGTTCGTCATCCC<br>ATCAGGACACTCGCGCAGTCCTTATTTCTGCCAGCCGCACCGCCTCGTCAGG<br>TCCTTCTGTGTCTTCGGCATATTCTGAAGATTTGCCGCATTTCGAATGCGCT<br>TTTTTCCTTCCGCATCATCATTCTGTCACGGAAACCGATTACCTCGCACTCC<br>TGACCTTGCGGGAAGCGGGCGAGTCACGCCAAATAGCTTCTCATCCCTGGTT<br>TCCCATGACACTGAAGATACT |
| **RBS *B0034***<br>Golden Gate overhangs | TACTAGAGAAAGAGGAGAAATAATCAATG |
| **CDS mScarlet**<br>Golden Gate overhangs<br>**ATG** | AATGGTTTCTAAAGGTGAAGCAGTGATCAAAGAATTTATGCGCTTCAAAGTT<br>CACATGGAAGGTTCTATGAACGGCCACGAATTTGAAATTGAAGGTGAAGGCG<br>AAGGTCGTCCATACGAAGGTACTCAAACCGCAAAACTGAAAGTTACCAAAGG<br>TGGTCCACTGCCATTTTCTTGGGATATTCTGTCTCCACAATTTATGTACGGT<br>TCTCGTGCATTTATCAAACACCCAGCAGATATTCCAGACTACTACAAACAAT<br>CTTTTCCGGAAGGTTTCAAATGGGAACGTGTTATGAATTTTGAAGATGGTGG<br>TGCAGTTACGGTTACCCAAGATACCTCTCTGGAAGATGGTACTCTGATCTAC<br>AAAGTTAAACTGCGTGGTACTAACTTTCCACCAGATGGTCCAGTTATGCAGA<br>AAAAAACCATGGGTTGGGAAGCATCTACCGAACGTCTGTACCCAGAAGATGG<br>CGTTCTGAAAGGTGATATCAAAATGGCACTGCGTCTGAAAGATGGCGGTCGT<br>TACCTGGCAGATTTCAAAACCACCTACAAAGCGAAAAAACCAGTTCAAATGC<br>CAGGTGCATACAACGTTGATCGTAAACTGGATATTACCAGCCACAACGAAGA<br>TTACACCGTTGTTGAACAATACGAACGTTCTGAAGGCCGTCACTCTACCGGT<br>GGTATGGATGAACTGTACAAAGCTT |
| **Terminator TB1002**<br>Golden Gate overhangs<br>***STOP* codon** | GCTTAACGCAAAAAACCCCGCTTCGGCGGGGTTTTTTCGCCGCT |
| **3' Connector 3C2SR**<br>Golden Gate overhangs | CGCTAGTAGGAGACGAGCT |
| **Origin of replication (*oriV*) ORSF1010**<br>Golden Gate overhangs | AGCTTTCTGAAAGCGACCAGGTGCTCGGCGTGGCAAGACTCGCAGCGAACCC<br>GTAGAAAGCCATGCTCCAGCCGCCCGCATTGGAGAAATTCTTCAAATTCCCG<br>TTGCACATAGCCCGGCAATTCCTTTCCCTGCTCTGCCATAAGCGCAGCGAAT<br>GCCGGGTAATACTCGTCAACGATCTGATAGAGAAGGGTTTGCTCGGGTCGGT<br>GGCTCTGGTAACGACCAGTATCCCGATCCCGGCTGGCCGTCCTGGCCGCCAC<br>ATGAGGCATGTTCCGCGTCCTTGCAATACTGTGTTTACATACAGTCTATCGC<br>TTAGCGGAAAGTTCTTTTACCCTCAGCCGAAATGCCTGCCGTTGCTAGACAT<br>TGCCAGCCAGTGCCCGTCACTCCCGTACTAACTGTCACGAACCCCTGCAATA<br>ACTGTCACGCCCCCCTGCAATAACTGTCACGAACCCCTGCAATAACTGTCAC<br>GCCCCCAAACCTGCAAACCCAGCAGGGGCGGGGGCTGGCGGGGTGTTGGAAA<br>AATCCATCCATGATTATCTAAGAATAATCCACTAGGCGCGGTTATCAGCGCC<br>CTTGTGGGGCGCTGCTGCCCTTGCCCAATATGCCCGGCCAGAGGCCGGATAG<br>CTGGTCTATTCGCTGCGCTAGGCTACACACCGCCCCACCGCTGCGCGGCAGG<br>GGGAAAGGCGGGCAAAGCCCGCTAAACCCCACACCAAACCCCGCAGAAATAC<br>GCTGGAGCGCTTTTAGCCGCTTTAGCGGCCTTTCCCCCTACCCGAAGGGTGG<br>GGGCGCGTGTGCAGCCCCGCAGGGCCTGTCTCGGTCGATCATTCAGCCCGGC<br>TCATCCTTCTGGCGTGGCGGCAGACCGAACAAGGCGCGGTCGTGGTCGCGTT<br>CAAGGTACGCATCCATTGCCGCCATGAGCCGATCCTCCGGCCACTCGCTGCT<br>GTTCACCTTGGCCAAAATCATGGCCCCCACCAGCACCTTGCGCCTTGTTTCG<br>TTCTTGCGCTCTTGCTGCTGTTCCCTTGCCCGCACCCGCTGAATTTCGGCAT<br>TGATTCGCGCTCGTTGTTCTTCGAGCTTGGCCAGCCGATCCGCCGCCTTGTT<br>GCTCCCCTTAACCATCTTGACACCCCATTGTTAATGTGCTGTCTCGTAGGCT<br>ATCATGGAGGCACAGCGGCGGCAATCCCGACCCTACTTTGTAGGGGAGGGCG<br>CACTTACCGGTTTCTCTTCGAGAAACTGGCCTAACGGCCACCCTTCGGGCGG<br>TGCGCTCTCCGAGGGCCATTGCATGGAGCCGAAAAGCAAAAGCAACAGCGAG<br>GCAGCATGGCGATTTATCACCTTACGGCGAAAACCGGCAGCAGGTCGGGCGG<br>CCAATCGGCCAGGGCCAAGGCCGACTACATCCAGCGCGAAGGCAAGTATGCC<br>CGCGACATGGATGAAGTCTTGCACGCCGAATCCGGGCACATGCCGGAGTTCG<br>TCGAGCGGCCCGCCGACTACTGGATGCTGCCGACCTGTATGAACGCGCCAA |

TGGGCGGCTGTTCAAGGAGGTCGAATTTGCCCTGCCGGTCGAGCTGACCCTC
GACCAGCAGAAGGCGCTGGCGTCCGAGTTCGCCCAGCACCTGACCGGTGCCG
AGCGCCTGCCGTATACGCTGGCCATCCATGCCGGTGGCGGCGAGAACCCGCA
CTGCCACCTGATGATCTCCGAGCGGATCAATGACGGCATCGAGCGGCCCGCC
GCTCAGTGGTTCAAGCGGTACAACGGCAAGACCCCGGAGAAGGGCGGGGCAC
AGAAGACCGAAGCGCTCAAGCCCAAGGCATGGCTTGAGCAGACCCGCGAGGC
ATGGGCCGACCATGCCAACCGGGCATTAGAGCGGGCTGGCCACGACGCCCGC
ATTGACCACAGAACACTTGAGGCGCAGGGCATCGAGCGCCTGCCCGGTGTTC
ACCTGGGGCCGAACGTGGTGGAGATGGAAGGCCGGGGCATCCGCACCGACCG
GGCAGACGTGGCCCTGAACATCGACACCGCCAACGCCCAGATCATCGACTTA
CAGGAATACCGGGAGGCAATAGACCATGAACGCAATCGACAGAGTGAAGAAA
TCCAGAGGCATCAACGAGTTAGCGGAGCAGATCGAACCGCTGGCCCAGAGCA
TGGCGACACTGGCCGACGAAGCCCGGCAGGTCATGAGCCAGACCAAGCAGGC
CAGCGAGGCGCAGGCGGCGGAGTGGCTGAAAGCCCAGCGCCAGACAGGGGCG
GCATGGGTGGAGCTGGCCAAAGAGTTGCGGGAGGTAGCCGCCGAGGTGAGCA
GCGCCGCGCAGAGCGCCCGGAGCGCGTCGCGGGGGTGGCACTGGAAGCTATG
GCTAACCGTGATGCTGGCTTCCATGATGCCTACGGTGGTGCTGCTGATCGCA
TCGTTGCTCTTGCTCGACCTGACGCCACTGACAACCGAGGACGGCTCGATCT
GGCTGCGCTTGGTGGCCCGATGAAGAACGACAGGACTTTGCAGGCCATAGGC
CGACAGCTCAAGGCCATGGGCTGTGAGCGCTTCGATATCGGCGTCAGGGACG
CCACCACCGGCCAGATGATGAACCGGGAATGGTCAGCCGCCGAAGTGCTCCA
GAACACGCCATGGCTCAAGCGGATGAATGCCCAGGGCAATGACGTGTATATC
AGGCCCGCCGAGCAGGAGCGGCATGGTCTGGTGCTGGTGGACGACCTCAGCG
AGTTTGACCTGGATGACATGAAAGCCGAGGGCCGGGAGCCTGCCCTGGTAGT
GGAAACCAGCCCGAAGAACTATCAGGCATGGGTCAAGGTGGCCGACGCCGCA
GGCGGTGAACTTCGGGGGCAGATTGCCCGGACGCTGGCCAGCGAGTACGACG
CCGACCCGGCCAGCGCCGACAGCCGCCACTATGGCCGCTTGGCGGGCTTCAC
CAACCGCAAGGACAAGCACACCACCCGCGCCGGTTATCAGCCGTGGGTGCTG
CTGCGTGAATCCAAGGGCAAGACCGCCACCGCTGGCCCGGCGCTGGTGCAGC
AGGCTGGCCAGCAGATCGAGCAGGCCCAGCGGCAGCAGGAGAAGGCCCGCAG
GCTGGCCAGCCTCGAACTGCCCGAGCGGCAGCTTAGCCGCCACCGGCGCACG
GCGCTGGACGAGTACCGCAGCGAGATGGCCGGGCTGGTCAAGCGCTTCGGTG
ATGACCTCAGCAAGTGCGACTTTATCGCCGCGCAGAAGCTGGCCAGCCGGGG
CCGCAGTGCCGAGGAAATCGGCAAGGCCATGGCCGAGGCCAGCCCAGCGCTG
GCAGAGCGCAAGCCCGGCCACGAAGCGGATTACATCGAGCGCACCGTCAGCA
AGGTCATGGGTCTGCCCAGCGTCCAGCTTGCGCGGGCCGAGCTGGCACGGGC
ACCGGCACCCCGCCAGCGAGGCATGGACAGGGGCGGGCCAGATTTCAGCATG
TAGTGCTTGCGTTGGTACTCACGCCTGTTATACTATGAGTACTCACGCACAG
AAGGGGGTTTTATGGAATACGAAAAAAGCGCTTCAGGGTCGGTCTACCTGAT
CAAAAGTGACAAGGGCTATTGGTTGCCCGGTGGCTTTGGTTATACGTCAAAC
AAGGCCGAGGCTGGCCGCTTTTCAGTCGCTGATATGGCCAGCCTTAACCTTG
ACGGCTGCACCTTGTCCTTGTTCCGCGAAGACAAGCCTTTCGGCCCCGGCAA
GTTTCTCGGTGACTGATATGAAAGACCAAAAGGACAAGCAGACCGGCGACCT
GCTGGCCAGCCCTGACGCTGTACGCCAAGCGCGATATGCCGAGCGCATGAAG
GCCAAAGGGATGCGTCAGCGCAAGTTCTGGCTGACCGACGACGAATACGAGG
CGCTGCGCGAGTGCCTGGAAGAACTCAGAGCGGCGCAGGGCGGGGGTAGTGA
CCCCGCCAGCGCCTAACCACCAACTGCCTGCAAAGGAGGCAATCAATGGCTA
CCCATAAGCCTATCAATATTCTGGAGGCGTTCGCAGCAGCGCCGCCACCGCT
GGACTACGTTTTGCCCAACATGGTGGCCGGTACGGTCGGGGCGCTGGTGTCG
CCCGGTGGTGCCGGTAAATCCATGCTGGCCCTGCAACTGGCCGCACAGATTG
CAGGCGGGCCGGATCTGCTGGAGGTGGGCGAACTGCCCACCGGCCCGGTGAT
CTACCTGCCCGCCGAAGACCCGCCCACCGCCATTCATCACCGCCTGCACGCC
CTTGGGGCGCACCTCAGCGCCGAGGAACGGCAAGCCGTGGCTGACGGCCTGC
TGATCCAGCCGCTGATCGGCAGCCTGCCCAACATCATGGCCCCGGAGTGGTT
CGACGGCCTCAAGCGCGCCGCCGAGGGCCGCCGCCTGATGGTGCTGGACACG
CTGCGCCGGTTCCACATCGAGGAAGAAAACGCCAGCGGCCCCATGGCCCAGG
TCATCGGTCGCATGGAGGCCATCGCCGCCGATACCGGGTGCTCTATCGTGTT
CCTGCACCATGCCAGCAAGGGCGCGGCCATGATGGGCGCAGGCGACCAGCAG
CAGGCCAGCCGGGGCAGCTCGGTACTGGTCGATAACATCCGCTGGCAGTCCT
ACCTGTCGAGCATGACCAGCGCCGAGGCCGAGGAATGGGGTGTGGACGACGA
CCAGCGCCGGTTCTTCGTCCGCTTCGGTGTGAGCAAGGCCAACTATGGCGCA

| | |
|---|---|
| | **CCGTTCGCTGATCGGTGGTTCAGGCGGCATGACGGCGGGGTGCTCAAGCCCG**<br>**CCGTGCTGGAGAGGCAGCGCAAGAGCAAGGGGGTGCCCCGTGGTGAAGCCTA**<br>**AGAACAAGCACAGCCTCAGCCACGTCCGGCACGACCCGGCGCACTGTCTGGC**<br>**CCCCGGCCTGTTCCGTGCCCTCAAGCGGGGCGAGCGCAAGCGCAGCAAGCTG**<br>**GACGTGACGTATGACTACGGCGACGGCAAGCGGATCGAGTTCAGCGGCCCGG**<br>**AGCCGCTGGGCGCTGATGATCTGCGCATCCTGCAAGGGCTGGTGGCCATGGC**<br>**TGGGCCTAATGGCCTAGTGCTTGGCCCGGAACCCAAGACCGAAGGCGGACGG**<br>**CAGCTCCGGCTGTTCCTGGAACCCAAGTGGGAGGCCGTCACCGCTGATGCCA**<br>**TGGTGGTCAAAGGTAGCTATCGGGCGCTGGCAAAGGAAATCGGGGCAGAGGT**<br>**CGATAGTGGTGGGGCGCTCAAGCACATACAGGACTGCATCGAGCGCCTTTGG**<br>**AAGGTATCCATCATCGCCCAGAATGGCCGCAAGCGGCAGGGGTTTCGGCTGC**<br>**TGTCGGAGTACGCCAGCGACGAGGCGGACGGGCGCCTGTACGTGGCCCTGAA**<br>**CCCCTTGATCGCGCAGGCCGTCATGGGTGGCGGCCAGCATGTGCGCATCAGC**<br>**ATGGACGAGGTGCGGGCGCTGGACAGCGAAACCGCCCGCCTGCTGCACCAGC**<br>**GGCTGTGTGGCTGGATCGACCCCGGCAAAACCGGCAAGGCTTCCATAGATAC**<br>**CTTGTGCGGCTATGTCTGGCCGTCAGAGGCCAGTGGTTCGACCATGCGCAAG**<br>**CGCCGCCAGCGGGTGCGCGAGGCGTTGCCGGAGCTGGTCGCGCTGGGCTGGA**<br>**CGGTAACCGAGTTCGCGGCGGGCAAGTACGACATCACCCGGCCCAAGGCGGC**<br>**AGGCTGACCCCCCCCACTCTATTGTAAACAAGACATTTTTATCTTTTATATT**<br>**CAATGGCTTATTTTCCTGCTAATTGGTAATACCATGAAAAATACCATGCTCA**<br>**GAAAAGGCTTAACAATATTTTGAAAAATTGCCTACTGAGCGCTGCCGCACAG**<br>**CTCCATAGGCCGCTTTCCTGGCTTTGCTTCCAGATGTATGCTCTTCTGCTCC**<br>**CGAACGCCAGCAAGACGTAGCCCAGCGCGTCGGCCAGCTTGCAATTCGCGCT**<br>**AACTTACATTAATTGCGTTGCGC**TGCT |
| **Tetracycline Resistance cassette**<br>Golden Gate overhangs | TGCTGAGTCACTAAGGGCTAACTAACTAATTACGTAGCAATCAACTCACTGG<br>CTCACCTTCACGGGTGGGCCTTTCTTCGGCACGGGCAAATTGCTGAATATTC<br>CTTTTCTTAGACGTCAGGTGGCAAACGCGAAGTAATCTTTTCGGTTTTAAAG<br>AAAAAGGGCAGGGTGGTGACACCTTGCCCGTTTTTTTGCCGGAGCTAGTATT<br>A**TTAGGTCGAGGTGGCCCGGCTCCATGCACCGCGACGCAACGCGGGGAGGCA**<br>**GACAAGGTAAAGGGCGGCACCTACAATCCATGCCAACCCGTTCCATGTGCTC**<br>**GCCGAGGCGGCATAAATCGCCGTGACGATCAGCGGTCCAATGATCGAAGTGA**<br>**GGCTGGTAAGAGCCGCGAGCGACCCTTGAAGCTGTCCCTGATGGTCGTCATC**<br>**TACTTGGCGGGACAGCATGGCCTGCAACGCGGGCATACCGATGCCGCCGGAA**<br>**GCGAGAAGAATCATAATGGGGAAGGCCATCCAGCCGCGCGTCGCGAACGCCA**<br>**GCAAGACGTAGCCCAGCGCGTCGGCCGCCATGCCCGCGATAATGGCCTGCTT**<br>**CTCGCCGAAACGTTTGGTGGCGGGGCCCGTGACGAAGGCTTGAGCGAGGGCG**<br>**TGCAAGATTCCGAATACCGCAAGCGACAGGCCGATCATCGTCGCGCTCCAGC**<br>**GAAAGCGGTCCTCGCCGAAAATGACCCAGAGCGCTGCCGGAACCTGTCCTAC**<br>**GAGTTGCATGATAAAGAAAACAGTCATAAGTGCGGCGACGATAGTCATGCCC**<br>**CGCGCCCACCGGAAGGAGCTGACTGGATTGAAGGCACGCAAGGGCATCGGAC**<br>**GGCGCTCTCCCTTATGCGATTCCTGCATAAGGAAGCAGCCCAGGAGGAGGTT**<br>**GAGGCCGTTGAGCACCGCCGCCGCAAGGAATGGTGCATGCAAGGAGATGGCA**<br>**CCCAACAGTCCCCCGGCCACGGGGCCTGCCACCATACCCACGCCGAAACAAG**<br>**CGCTCATGAGCCCGAAGTGGCGAGCCCGATCTTCCCCATCGGTGATGTCGGC**<br>**GATATAGGCACCAGCAACCGCACCTGTGGCACCCGTGATGCCCGCCACGATG**<br>**CGTCCGGCGTAGAGAATCCACAGGACGGGTGTGGTCGCCATGATCGCGTAGT**<br>**CGATAGTGGCTCCAAGGAGCGAAGCGAGCAGGACTGGACGGCGGCCAAAGCG**<br>**GTCGGACAGGGCTCCGAGAACGGGTGCGCAAAGAAATTGCATCAACGCATAG**<br>**AGCGCAAGCAGCACGCCATAGTGACTGGCAATACTGTCGGAATGGACGATAT**<br>**CCCGCAAGAGGCCCGGCAGTACCGGCATAACCAAGCCTATGCCTACAGCGTC**<br>**CAGGGTGACGGTGCCGAGAATGACGATGAGCGCATTGTTAGATTTCAT**ACAC<br>GGTGCCTGACTGCGTTAGCAATTTAACTGTGATAAACTACCGCATTAAAGCT<br>TATCGATGATAAGCTGTCAAACATGAGAATCTCCTCTAGTAAACACCCCTTG<br>TATTACTGTTTATGTAAGCAGACAGTTTTATTGTTCATGATGATATATTTTT<br>ATCTTGTGCAATGTAACATCAGAGATTTTGAGACACAACGTGGCTTTGTTGA<br>ATAAATCGAACTTTTGCTGAGTTGAAGGATCAGATCACGCATCTTAACA |
| pL1_FP*phnG* plasmid<br>**Tetracycline Resistance cassette**<br>**5' Connector 5C5CSR** | TGCTGAGTCACTAAGGGCTAACTAACTAATTACGTAGCAATCAACTCACTGG<br>CTCACCTTCACGGGTGGGCCTTTCTTCGGCACGGGCAAATTGCTGAATATTC<br>CTTTTCTTAGACGTCAGGTGGCAAACGCGAAGTAATCTTTTCGGTTTTAAAG<br>AAAAAGGGCAGGGTGGTGACACCTTGCCCGTTTTTTTGCCGGAGCTAGTATT<br>A**TTAGGTCGAGGTGGCCCGGCTCCATGCACCGCGACGCAACGCGGGGAGGCA** |

| | |
|---|---|
| Promoter $P_{phnG}$<br>RBS B0034<br>CDS mScarlet<br>Terminator TB1002<br>3' Connector 3C2SR<br>**Origin of replication (ORI) ORSF1010**<br>Golden Gate overhangs | GACAAGGTAAAGGGCGGCACCTACAATCCATGCCAACCCGTTCCATGTGCTC<br>GCCGAGGCGGCATAAATCGCCGTGACGATCAGCGGTCCAATGATCGAAGTGA<br>GGCTGGTAAGAGCCGCGAGCGACCCTTGAAGCTGTCCCTGATGGTCGTCATC<br>TACTTGGCGGGACAGCATGGCCTGCAACGCGGGCATACCGATGCCGCCGGAA<br>GCGAGAAGAATCATAATGGGGAAGGCCATCCAGCCGCGCGTCGCGAACGCCA<br>GCAAGACGTAGCCCAGCGCGTCGGCCGCCATGCCCGCGATAATGGCCTGCTT<br>CTCGCCGAAACGTTTGGTGGCGGGGCCCGTGACGAAGGCTTGAGCGAGGGCG<br>TGCAAGATTCCGAATACCGCAAGCGACAGGCCGATCATCGTCGCGCTCCAGC<br>GAAAGCGGTCCTCGCCGAAAATGACCCAGAGCGCTGCCGGAACCTGTCCTAC<br>GAGTTGCATGATAAAGAAAACAGTCATAAGTGCGGCGACGATAGTCATGCCC<br>CGCGCCCACCGGAAGGAGCTGACTGGATTGAAGGCACGCAAGGGCATCGGAC<br>GGCGCTCTCCCTTATGCGATTCCTGCATAAGGAAGCAGCCCAGGAGGAGGTT<br>GAGGCCGTTGAGCACCGCCGCCGCAAGGAATGGTGCATGCAAGGAGATGGCA<br>CCCAACAGTCCCCCGGCCACGGGGCCTGCCACCATACCCACGCCGAAACAAG<br>CGCTCATGAGCCCGAAGTGGCGAGCCCGATCTTCCCCATCGGTGATGTCGGC<br>GATATAGGCACCAGCAACCGCACCTGTGGCACCCGTGATGCCCGCCACGATG<br>CGTCCGGCGTAGAGAATCCACAGGACGGGTGTGGTCGCCATGATCGCGTAGT<br>CGATAGTGGCTCCAAGGAGCGAAGCGAGCAGGACTGGACGGCGGCCAAAGCG<br>GTCGGACAGGGCTCCGAGAACGGGTGCGCAAAGAAATTGCATCAACGCATAG<br>AGCGCAAGCAGCACGCCATAGTGACTGGCAATACTGTCGGAATGGACGATAT<br>CCCGCAAGAGGCCCGGCAGTACCGGCATAACCAAGCCTATGCCTACAGCGTC<br>CAGGGTGACGGTGCCGAGAATGACGATGAGCGCATTGTTAGATTTCATACAC<br>GGTGCCTGACTGCGTTAGCAATTTAACTGTGATAAACTACCGCATTAAAGCT<br>TATCGATGATAAGCTGTCAAACATGAGAATCTCCTCTAGTAAACACCCCTTG<br>TATTACTGTTTATGTAAGCAGACAGTTTTATTGTTCATGATGATATATTTTT<br>ATCTTGTGCAATGTAACATCAGAGATTTTGAGACACAACGTGGCTTTGTTGA<br>ATAAATCGAACTTTTGCTGAGTTGAAGGATCAGATCACGCATCTTAACACGT<br>CTCGAGCGGGAGTCAATTTTCCACCGTGAATTCGACCGTATCGGCCGAAAAA<br>CGGCTGATGGCGTATTGCACCGGCACGCCATCCATATCGGCGTTCAGCGCCT<br>GGGTGACGAGCAGGATCGCGCCCGGCGACAATTCGAGATCGGCCAGATCCTG<br>CGTATCCGCATGGCTGGCGGAGATGACGGTGGAAATGCGCACGTAATCCGCC<br>ACGCCGAGCATCTGGAAAGCGGCCGTGATCGAGCCGCTTTTCTGATAGGCCT<br>CGCCTATTCCGGCAAAACGGTCGGCCGGGAACCATGTGGTCGAGCGGGATAC<br>AGGGCGGTTATCCGCCTTGCGCAGCGTTTCGAGGCGCACGAGAGGCGCGCCT<br>GATGGCACTTGCAGTCGCGCCGCCAGATCGGCATTGGCCGGCTCGGTGGTAT<br>GGGACAGAAGCAGCGCCTCCATTTCCTTCACCTGATCGCCAATGCCCTCCGT<br>AAATCGCGTGCGCCGTGAAATCGGAAAACTCAGGCGCTCCTTGCGGGCGATC<br>ATCGTGCCACGCCCCTGCATTGGCTCGAGAATGCCTTCACTGGCCAATGCGG<br>CAATGGCGCTTCTCACAGTATGGCGGTTGACGCCGAATTTTTCAGCCAGCGC<br>CATTTCCGGCGGCAACATGCCGGTTTCGTCGTGATCGCCTGCCGCGATCTCG<br>CCTCTGATCCGGTCGGCGATCTGCCGCCAGAGCGCCACGCCGTTCTTTCTTT<br>TCATCGCTGCCGCAGGGCTCATGTCACACGCTTGTCATTCATCGACGTTAAT<br>TGTAGCTTATATTGTATAGTTGTCTATATCAATAGACATTAGAGGTCAAGCG<br>TACTAGAGAAAGAGGAGAAATAATCAATGGTTTCTAAAGGTGAAGCAGTGAT<br>CAAAGAATTTATGCGCTTCAAAGTTCACATGGAAGGTTCTATGAACGGCCAC<br>GAATTTGAAATTGAAGGTGAAGGCGAAGGTCGTCCATACGAAGGTACTCAAA<br>CCGCAAAACTGAAAGTTACCAAAGGTGGTCCACTGCCATTTTCTTGGGATAT<br>TCTGTCTCCACAATTTATGTACGGTTCTCGTGCATTTATCAAACACCCAGCA<br>GATATTCCAGACTACTACAAACAATCTTTTCCGGAAGGTTTCAAATGGGAAC<br>GTGTTATGAATTTTGAAGATGGTGGTGCAGTTACGGTTACCCAAGATACCTC<br>TCTGGAAGATGGTACTCTGATCTACAAAGTTAAACTGCGTGGTACTAACTTT<br>CCACCAGATGGTCCAGTTATGCAGAAAAAAACCATGGGTTGGGAAGCATCTA<br>CCGAACGTCTGTACCCAGAAGATGGCGTTCTGAAAGGTGATATCAAAATGGC<br>ACTGCGTCTGAAAGATGGCGGTCGTTACCTGGCAGATTTCAAAACCACCTAC<br>AAAGCGAAAAAACCAGTTCAAATGCCAGGTGCATACAACGTTGATCGTAAAC<br>TGGATATTACCAGCCACAACGAAGATTACACCGTTGTTGAACAATACGAACG<br>TTCTGAAGGCCGTCACTCTACCGGTGGTATGGATGAACTGTACAAAGCTTAA<br>CGCAAAAAACCCCGCTTCGGCGGGTTTTTTCGCCGCTAGTAGGAGACGAGC<br>TTTCTGAAAGCGACCAGGTGCTCGGCGTGGCAAGACTCGCAGCGAACCCGTA<br>GAAAGCCATGCTCCAGCCGCCCGCATTGGAGAAATTCTTCAAATTCCCGTTG<br>CACATAGCCCGGCAATTCCTTTCCCTGCTCTGCCATAAGCGCAGCGAATGCC |

GGGTAATACTCGTCAACGATCTGATAGAGAAGGGTTTGCTCGGGTCGGTGGC
TCTGGTAACGACCAGTATCCCGATCCCGGCTGGCCGTCCTGGCCGCCACATG
AGGCATGTTCCGCGTCCTTGCAATACTGTGTTTACATACAGTCTATCGCTTA
GCGGAAAGTTCTTTTACCCTCAGCCGAAATGCCTGCCGTTGCTAGACATTGC
CAGCCAGTGCCCGTCACTCCCGTACTAACTGTCACGAACCCCTGCAATAACT
GTCACGCCCCCCTGCAATAACTGTCACGAACCCCTGCAATAACTGTCACGCC
CCCAAACCTGCAAACCCAGCAGGGGCGGGGGCTGGCGGGGTGTTGGAAAAAT
CCATCCATGATTATCTAAGAATAATCCACTAGGCGCGGTTATCAGCGCCCTT
GTGGGGCGCTGCTGCCCTTGCCCAATATGCCCGGCCAGAGGCCGGATAGCTG
GTCTATTCGCTGCGCTAGGCTACACACCGCCCCACCGCTGCGCGGCAGGGGG
AAAGGCGGGCAAAGCCCGCTAAACCCCACACCAAACCCCGCAGAAATACGCT
GGAGCGCTTTTAGCCGCTTTAGCGGCCTTTCCCCCTACCCGAAGGGTGGGGG
CGCGTGTGCAGCCCCGCAGGGCCTGTCTCGGTCGATCATTCAGCCCGGCTCA
TCCTTCTGGCGTGGCGGCAGACCGAACAAGGCGCGGTCGTGGTCGCGTTCAA
GGTACGCATCCATTGCCGCCATGAGCCGATCCTCCGGCCACTCGCTGCTGTT
CACCTTGGCCAAAATCATGGCCCCCACCAGCACCTTGCGCCTTGTTTCGTTC
TTGCGCTCTTGCTGCTGTTCCCTTGCCCGCACCCGCTGAATTTCGGCATTGA
TTCGCGCTCGTTGTTCTTCGAGCTTGGCCAGCCGATCCGCCGCCTTGTTGCT
CCCCTTAACCATCTTGACACCCCATTGTTAATGTGCTGTCTCGTAGGCTATC
ATGGAGGCACAGCGGCGGCAATCCCGACCCTACTTTGTAGGGGAGGGCGCAC
TTACCGGTTTCTCTTCGAGAAACTGGCCTAACGGCCACCCTTCGGGCGGTGC
GCTCTCCGAGGGCCATTGCATGGAGCCGAAAAGCAAAAGCAACAGCGAGGCA
GCATGGCGATTTATCACCTTACGGCGAAAACCGGCAGCAGGTCGGGCGGCCA
ATCGGCCAGGGCCAAGGCCGACTACATCCAGCGCGAAGGCAAGTATGCCCGC
GACATGGATGAAGTCTTGCACGCCGAATCCGGGCACATGCCGGAGTTCGTCG
AGCGGCCCGCCGACTACTGGGATGCTGCCGACCTGTATGAACGCGCCAATGG
GCGGCTGTTCAAGGAGGTCGAATTTGCCCTGCCGGTCGAGCTGACCCTCGAC
CAGCAGAAGGCGCTGGCGTCCGAGTTCGCCCAGCACCTGACCGGTGCCGAGC
GCCTGCCGTATACGCTGGCCATCCATGCCGGTGGCGGCGAGAACCCGCACTG
CCACCTGATGATCTCCGAGCGGATCAATGACGGCATCGAGCGGCCCGCCGCT
CAGTGGTTCAAGCGGTACAACGGCAAGACCCCGGAGAAGGGCGGGGCACAGA
AGACCGAAGCGCTCAAGCCCAAGGCATGGCTTGAGCAGACCCGCGAGGCATG
GGCCGACCATGCCAACCGGGCATTAGAGCGGGCTGGCCACGACGCCCGCATT
GACCACAGAACACTTGAGGCGCAGGGCATCGAGCGCCTGCCCGGTGTTCACC
TGGGGCCGAACGTGGTGGAGATGGAAGGCCGGGGCATCCGCACCGACCGGGC
AGACGTGGCCCTGAACATCGACACCGCCAACGCCCAGATCATCGACTTACAG
GAATACCGGGAGGCAATAGACCATGAACGCAATCGACAGAGTGAAGAAATCC
AGAGGCATCAACGAGTTAGCGGAGCAGATCGAACCGCTGGCCCAGAGCATGG
CGACACTGGCCGACGAAGCCCGGCAGGTCATGAGCCAGACCAAGCAGGCCAG
CGAGGCGCAGGCGGCGGAGTGGCTGAAAGCCCAGCGCCAGACAGGGGCGGCA
TGGGTGGAGCTGGCCAAAGAGTTGCGGGAGGTAGCCGCCGAGGTGAGCAGCG
CCGCGCAGAGCGCCCGGAGCGCGTCGCGGGGGTGGCACTGGAAGCTATGGCT
AACCGTGATGCTGGCTTCCATGATGCCTACGGTGGTGCTGCTGATCGCATCG
TTGCTCTTGCTCGACCTGACGCCACTGACAACCGAGGACGGCTCGATCTGGC
TGCGCTTGGTGGCCCGATGAAGAACGACAGGACTTTGCAGGCCATAGGCCGA
CAGCTCAAGGCCATGGGCTGTGAGCGCTTCGATATCGGCGTCAGGGACGCCA
CCACCGGCCAGATGATGAACCGGGAATGGTCAGCCGCCGAAGTGCTCCAGAA
CACGCCATGGCTCAAGCGGATGAATGCCCAGGGCAATGACGTGTATATCAGG
CCCGCCGAGCAGGAGCGGCATGGTCTGGTGCTGGTGGACGACCTCAGCGAGT
TTGACCTGGATGACATGAAAGCCGAGGGCCGGGAGCCTGCCCTGGTAGTGGA
AACCAGCCCGAAGAACTATCAGGCATGGGTCAAGGTGGCCGACGCCGCAGGC
GGTGAACTTCGGGGGCAGATTGCCCGGACGCTGGCCAGCGAGTACGACGCCG
ACCCGGCCAGCGCCGACAGCCGCCACTATGGCCGCTTGGCGGGCTTCACCAA
CCGCAAGGACAAGCACACCACCCGCGCCGGTTATCAGCCGTGGGTGCTGCTG
CGTGAATCCAAGGGCAAGACCGCCACCGCTGGCCCGGCGCTGGTGCAGCAGG
CTGGCCAGCAGATCGAGCAGGCCCAGCGGCAGCAGGAGAAGGCCCGCAGGCT
GGCCAGCCTCGAACTGCCCGAGCGGCAGCTTAGCCGCCACCGGCGCACGGCG
CTGGACGAGTACCGCAGCGAGATGGCCGGGCTGGTCAAGCGCTTCGGTGATG
ACCTCAGCAAGTGCGACTTTATCGCCGCGCAGAAGCTGGCCAGCCGGGGCCG
CAGTGCCGAGGAAATCGGCAAGGCCATGGCCGAGGCCAGCCCAGCGCTGGCA
GAGCGCAAGCCCGGCCACGAAGCGGATTACATCGAGCGCACCGTCAGCAAGG

TCATGGGTCTGCCCAGCGTCCAGCTTGCGCGGGCCGAGCTGGCACGGGCACC
GGCACCCCGCCAGCGAGGCATGGACAGGGGCGGGCCAGATTTCAGCATGTAG
TGCTTGCGTTGGTACTCACGCCTGTTATACTATGAGTACTCACGCACAGAAG
GGGGTTTTATGGAATACGAAAAAAGCGCTTCAGGGTCGGTCTACCTGATCAA
AAGTGACAAGGGCTATTGGTTGCCCGGTGGCTTTGGTTATACGTCAAACAAG
GCCGAGGCTGGCCGCTTTTCAGTCGCTGATATGGCCAGCCTTAACCTTGACG
GCTGCACCTTGTCCTTGTTCCGCGAAGACAAGCCTTTCGGCCCCGGCAAGTT
TCTCGGTGACTGATATGAAAGACCAAAAGGACAAGCAGACCGGCGACCTGCT
GGCCAGCCCTGACGCTGTACGCCAAGCGCGATATGCCGAGCGCATGAAGGCC
AAAGGGATGCGTCAGCGCAAGTTCTGGCTGACCGACGACGAATACGAGGCGC
TGCGCGAGTGCCTGGAAGAACTCAGAGCGGCGCAGGGCGGGGGTAGTGACCC
CGCCAGCGCCTAACCACCAACTGCCTGCAAAGGAGGCAATCAATGGCTACCC
ATAAGCCTATCAATATTCTGGAGGCGTTCGCAGCAGCGCCGCCACCGCTGGA
CTACGTTTTGCCCAACATGGTGGCCGGTACGGTCGGGGCGCTGGTGTCGCCC
GGTGGTGCCGGTAAATCCATGCTGGCCCTGCAACTGGCCGCACAGATTGCAG
GCGGGCCGGATCTGCTGGAGGTGGGCGAACTGCCCACCGGCCCGGTGATCTA
CCTGCCCGCCGAAGACCCGCCCACCGCCATTCATCACCGCCTGCACGCCCTT
GGGGCGCACCTCAGCGCCGAGGAACGGCAAGCCGTGGCTGACGGCCTGCTGA
TCCAGCCGCTGATCGGCAGCCTGCCCAACATCATGGCCCCGGAGTGGTTCGA
CGGCCTCAAGCGCGCCGCCGAGGGCCGCCGCCTGATGGTGCTGGACACGCTG
CGCCGGTTCCACATCGAGGAAGAAAACGCCAGCGGCCCCATGGCCCAGGTCA
TCGGTCGCATGGAGGCCATCGCCGCCGATACCGGGTGCTCTATCGTGTTCCT
GCACCATGCCAGCAAGGGCGCGGCCATGATGGGCGCAGGCGACCAGCAGCAG
GCCAGCCGGGGCAGCTCGGTACTGGTCGATAACATCCGCTGGCAGTCCTACC
TGTCGAGCATGACCAGCGCCGAGGCCGAGGAATGGGGTGTGGACGACGACCA
GCGCCGGTTCTTCGTCCGCTTCGGTGTGAGCAAGGCCAACTATGGCGCACCG
TTCGCTGATCGGTGGTTCAGGCGGCATGACGGCGGGGTGCTCAAGCCCGCCG
TGCTGGAGAGGCAGCGCAAGAGCAAGGGGGTGCCCCGTGGTGAAGCCTAAGA
ACAAGCACAGCCTCAGCCACGTCCGGCACGACCCGGCGCACTGTCTGGCCCC
CGGCCTGTTCCGTGCCCTCAAGCGGGGCGAGCGCAAGCGCAGCAAGCTGGAC
GTGACGTATGACTACGGCGACGGCAAGCGGATCGAGTTCAGCGGCCCGGAGC
CGCTGGGCGCTGATGATCTGCGCATCCTGCAAGGGCTGGTGGCCATGGCTGG
GCCTAATGGCCTAGTGCTTGGCCCGGAACCCAAGACCGAAGGCGGACGGCAG
CTCCGGCTGTTCCTGGAACCCAAGTGGGAGGCCGTCACCGCTGATGCCATGG
TGGTCAAAGGTAGCTATCGGGCGCTGGCAAAGGAAATCGGGGCAGAGGTCGA
TAGTGGTGGGGCGCTCAAGCACATACAGGACTGCATCGAGCGCCTTTGGAAG
GTATCCATCATCGCCCAGAATGGCCGCAAGCGGCAGGGGTTTCGGCTGCTGT
CGGAGTACGCCAGCGACGAGGCGGACGGGCGCCTGTACGTGGCCCTGAACCC
CTTGATCGCGCAGGCCGTCATGGGTGGCGGCCAGCATGTGCGCATCAGCATG
GACGAGGTGCGGGCGCTGGACAGCGAAACCGCCCGCCTGCTGCACCAGCGGC
TGTGTGGCTGGATCGACCCCGGCAAAACCGGCAAGGCTTCCATAGATACCTT
GTGCGGCTATGTCTGGCCGTCAGAGGCCAGTGGTTCGACCATGCGCAAGCGC
CGCCAGCGGGTGCGCGAGGCGTTGCCGGAGCTGGTCGCGCTGGGCTGGACGG
TAACCGAGTTCGCGGCGGGCAAGTACGACATCACCCGGCCCAAGGCGGCAGG
CTGACCCCCCCCACTCTATTGTAAACAAGACATTTTTATCTTTTATATTCAA
TGGCTTATTTTCCTGCTAATTGGTAATACCATGAAAAATACCATGCTCAGAA
AAGGCTTAACAATATTTTGAAAAATTGCCTACTGAGCGCTGCCGCACAGCTC
CATAGGCCGCTTTCCTGGCTTTGCTTCCAGATGTATGCTCTTCTGCTCCCGA
ACGCCAGCAAGACGTAGCCCAGCGCGTCGGCCAGCTTGCAATTCGCGCTAAC
TTACATTAATTGCGTTGCGC

**Table S2.** Vectors used for Golden Gate assembly of biosensor plasmids.

| Plasmid | Relevant characteristics | Reference |
|---|---|---|
| pV_01 | Golden Gate Level 0 plasmid, allows cloning of parts between BsaI sites | Stukenberg et al. [1] |
| pMC0_1_18 | Derivative of vector pV_01. Contains the 5' 5C5CSR linker | Stukenberg et al. [1] |
| pMC0_2_21_$P_{dummy}$ | Derivative of vector pV_01. Contains the non-coding promoter ($P_{dummy}$) | Stukenberg et al. [1] |
| pL0_2_ $P_{phnG}$ | Derivative of vector pV_01. Contains the $P_{phnG}$ promoter | This work |
| pL0_2_ $P_{phnJ}$ | Derivative of vector pV_01. Contains the $P_{phnJ}$ promoter | This work |
| pL0_2_ $P_{phnC}$ | Derivative of vector pV_01. Contains the $P_{phnC}$ promoter | This work |
| pL0_2_ $P_{phnD}$ | Derivative of vector pV_01. Contains the $P_{phnD}$ promoter | This work |
| pL0_2_ $P_{phnE2}$ | Derivative of vector pV_01. Contains the $P_{phnE2}$ promoter | This work |
| pL0_2_ $P_{duf1045}$ | Derivative of vector pV_01. Contains the $P_{duf1045}$ promoter | This work |
| pMC0_3_07 | Derivative of vector pV_01. Contains the *B0034* RBS | Stukenberg et al. [1] |
| pMC0_4_18_CDSmScarlet | Derivative of vector pV_01. Contains the *mScarlet* CDS | Stukenberg et al. [1] |
| pMC0_5_04 | Derivative of vector pV_01. Contains the TB002 terminator | Stukenberg et al. [1] |
| pMC0_6_09 | Derivative of vector pV_01. Contains the 3C2SR connector 3’ | Stukenberg et al. [1] |
| pMC0_7_05 | Derivative of vector pV_01. Contains the *ORI* (RSF1010) | Stukenberg et al. [1] |
| pMC0_8_11 | Derivative of vector pV_01. Contains the Resistance cassette for Tet | Stukenberg et al. [1] |

**Table S3.** Oligonucleotides used in this work**.**

| Name | Sequence | Use |
|---|---|---|
| FM152 | TTCGTCTCCCTCAAGTATGCGCGTTAGCCTTGTGAAAAAAGG | Cloning of $P_{phnJ}$ into L0 pV_01 plasmid |
| FM153 | AACGTCTCGCTCGGGAGCTCAGCGGCTGCCCTGCTCAC | |
| FM154 | AACGTCTCGCTCGGGAGGTCACACGCTTGTCATTCATC | Cloning of $P_{phnG}$ into L0 pV_01 plasmid |
| FM155 | TTCGTCTCCCTCAAGTACGCTTGACCTCTAATGTCTATTG | |
| FM156 | AACGTCTCGCTCGGGAGTCCTGACCCACCGTTCCGAAAG | Cloning of $P_{phnE2}$ into L0 pV_01 plasmid |
| FM157 | TTCGTCTCCCTCAAGTAGCCGCTTTCCTTTATTGGCAAAAACG | |
| FM158 | AACGTCTCGCTCGGGAGGGAACCGCTCAGCATCGTTTG | Cloning of $P_{phnD}$ into L0 pV_01 plasmid |
| FM159 | TTCGTCTCCCTCAAGTAGGACTTTCTCCTGTTTCCCGG | |
| FM160 | AACGTCTCGCTCGGGAGAATCTCACCTCTGGCAAGGCAAG | Cloning of $P_{phnC}$ into L0, pV_01 plasmid |
| FM161 | TTCGTCTCCCTCAAGTATTCCCCGTCCAATCTTCTTTTGAC | |
| FM162 | AACGTCTCGCTCGGGAGGCTTTGATGGGGTGGGATGAC | Cloning of $P_{duf1045}$ into L0 pV_01 plasmid |
| FM163 | TTCGTCTCCCTCAAGTATCTTCAGTGTCATGGGAAACCAGG | |
| FM171 | CATCAGAGATTTTGAGACACAACGTGGC | Cloning checking and sequencing of L1 plasmids |
| FM172 | GCGGCTGGAGCATGGCTTTCTA | |
| FM192 | GAATATCCCAAGAAAATGGCAGTGGACCAC | Sequencing of L1 plasmids, Scarlet CDS |
| FM241 | ATTAACGTCGATGACTCCCGCTCGAGACGTGTTAAG | Gibson assembly for pL1_$P_{phnG_\Delta B}$ |
| FM242 | TCGAGCGGGAGTCATCGACGTTAATTGTAGCTTATATTGTATAG | |
| FM243 | AGCGGGAGGTCACACGCTTGTCAT | Checking the pL1_$P_{phnG_\Delta B}$ cloning |
| FM244 | TCAAAAGTGACAAGGGCTATTGGTTGCC | Gibson assembly for pL1_$P_{phnG_\Delta B}$ |
| FM245 | GGCAACCAATAGCCCTTGTCACTTTTGA | |
| FM257 | GAATTCACGGTGGAAAATTGACTCCCGCTCGAGACGTGTTAAG | Gibson assembly for pL1_PhnF-$P_{phnG}$ |
| FM258 | GAGCGGGAGTCAATTTTCCACCGTGAATTCGAC | |
| FM259 | GCTGCCGCAGGGCTCATGTCACACGCTTGTCATTCATCGACG | Gibson assembly for pL1_PhnF-$P_{phnG}$ |
| FM260 | CAAGCGTGTGACATGAGCCCTGCGGCAGCGATG | |
| FM261 | ATATTGTATAGTTTAGAGGTCAAGCGTACTAGAGAAA | Gibson assembly for pL1_$P_{phnG_\Delta F}$ |
| FM262 | AGTACGCTTGACCTCTAAACTATACAATATAAGCTACAATTAAC | |
| FM263 | AGTTGTCTATATCAATAGACA | Checking the pL1_$P_{phnG_\Delta F}$ cloning |

**Table S4.** Whole-cell biosensor strains constructed in this work.

| Whole-cell biosensor strain | Characteristics |
|---|---|
| CHLDO·$P_{dummy}$ | *A. tumefaciens* CHLDO carrying plasmid pL1_$P_{dummy}$ |
| CHLDO·$P_{phnG}$ | *A. tumefaciens* CHLDO carrying plasmid pL1_$P_{phnG}$ |
| CHLDO·$P_{phnJ}$ | *A. tumefaciens* CHLDO carrying plasmid pL1_$P_{phnJ}$ |
| CHLDO·$P_{phnC}$ | *A. tumefaciens* CHLDO carrying plasmid pL1_$P_{phnC}$ |
| CHLDO·$P_{phnD}$ | *A. tumefaciens* CHLDO carrying plasmid pL1_$P_{phnD}$ |
| CHLDO·$P_{phnE2}$ | *A. tumefaciens* CHLDO carrying plasmid pL1_$P_{phnE2}$ |
| CHLDO·$P_{duf1045}$ | *A. tumefaciens* CHLDO carrying plasmid pL1_$P_{duf1045}$ |
| CHLDO·$P_{phnG\_\Delta B}$ | *A. tumefaciens* CHLDO carrying plasmid pL1_$P_{phnG_\Delta B}$ |
| CHLDO·$P_{phnG\_\Delta F}$ | *A. tumefaciens* CHLDO carrying plasmid pL1_$P_{phnG_\Delta F}$ |
| CHLDO·$FP_{phnG}$ | *A. tumefaciens* CHLDO carrying plasmid pL1_$FP_{phnG}$ |
| GV3101·$P_{dummy}$ | *A. tumefaciens* GV3101 carrying plasmid pGP1_$P_{dummy}$ |
| GV3101·$P_{phnG}$ | *A. tumefaciens* GV3101 carrying plasmid pL1_$P_{phnG}$ |
| GV3101·$P_{phnG\_sb}$ | *A. tumefaciens* GV3101 carrying plasmid pL1_$P_{phnG_\Delta B}$ |
| GV3101·$P_{phnG\_sf}$ | *A. tumefaciens* GV3101 carrying plasmid pL1_$P_{phnG_\Delta F}$ |
| GV3101·$FP_{phnG}$ | *A. tumefaciens* GV3101 carrying plasmid pL1_$FP_{phnG}$ |

**Table S5.** Summary of recent sensing technologies for GP detection.

| Detection technique | Characteristics | | | | | Reference |
|---|---|---|---|---|---|---|
| | Principle | Range | GP limit of detection | Equipment needed | Type of samples | |
| Electrochemical biosensor | A graphite-epoxy electrode (GE) modified with multiwalled carbon nanotubes and horseradish peroxidase inhibited by GP | N.R. | 1.32 pM | Potentiostat; galvanostat | Maize kernels | Cahuantzi-Muñoz et al. [2] |
| Enzymatic biosensor; potentiometric sensor | GP inhibits urease activity that is entrapped in agarose-guar gum matrix | 2.9 μM to 290 μM | 2.9 μM | Ion selective electrode to urea, urease immobilized membrane | Water samples | Vaghela et al. [3] |
| Enzymatic biosensor; amperometry | GP inhibition of tyrosinase enzyme. A carbon nano-onion/tyrosinase conjugate immobilized in a chitosan matrix on a screen-printed electrode | 0.015 to 10 μM | 6.5 nM | Potentiostat; galvanostat | Water and soil samples | Sok and Fragoso [4] |
| Enzymatic biosensor; colorimetric assay; ELISA | Surface immobilization of EPSPS *via* fusion to the hydrophobin Ccg2; GP inhibits EPSPS | 0.05-1 μM | 50 nM | Spectro-photometer, microplate reader at 630 nm | GP samples | Döring et al. [5] |
| Enzymatic biosensor; electrochemi-luminescence sensor | Based on double inhibition: GP inhibition of a peroxidase and formation a complex of GP-Cu(II) that prevents luminol-gold nanoparticles-L-cysteine-Cu(II) composites | 0.001 ~ 1.0 μM | 0.5 nM | Electro-luminescence analyzer | Water samples | Li et al. [6] |
| WCB, based on microalgae immobilization in an alginate matrix | GP inhibits algae growth; *Scenedesmus acutus* and *Pseudokirchneriella subcapitata* immobilized in alginate beads and tested as bioindicators | Naked eye: 14.5 to 290 μM; fluorescence: 29 to 118 μM | Naked eye: 88 μM for GP-based formulation | Naked eye and absorbance measurements and chlorophyll fluorescence | GBHs | Prudkin-Silva et al. [7] |
| WCB, based on cyanobacteria immobilization; hydrogel-gated organic field-effect transistor | *Anabaena flos-aquae* entrapped in a hydrogel grafted onto the platinum gate of the transistor; rates of oxygen produced or consumed by the cyanobacteria are affected by GP | N.R. | 10 μM | Photosynthesis monitored by electroreduction | GP chemical standard | Le Gall et al. [8] |
| Enzymatic biosensor; chronoamperometry | GP inhibits an acid phosphatase that is chemically immobilized to a carbon electrode modified with silver nanoparticles, reduced graphene oxide | Two linear ranges for GP from 0.2 μM to 2 μM and from 2.9 μM to 130 μM | 0.088 μM | Biosensor created manually and composed by three electrodes | GP-contaminated water and soil samples | Butmee et al. [9] |
| DNA-templated copper nanoparticles (DNA-CuNPs) | Fluorescence; GP strongly chelates $Cu^{2+}$ impeding the formation of DNA-CuNPs | 1 to 18 μM | 0.47 μM | Microplate readers with fluorescence excitation and detection | Agriculture samples (rice, maize, and wheat) | Fang et al. [10] |

| | | | | | | |
|---|---|---|---|---|---|---|
| Enzymatic biosensor | Fluorescence; GP affects the activity of the exonuclease I and the T5 exonuclease enzymes, slowing down enzymatic digestion | N.R. | 0.6 µM | Microplate readers with fluorescence measured at 545 nm (excitation at 495 nm) | GP samples | Berkal et al. [11] |
| Enzymatic biosensor; electrochemical sensor | Cyclic voltammetry and amperometry. GP inhibit of Horseradish peroxidase immobilized on a membrane doped with multiwalled carbon nanotubes | 0.5 to 59 µM | 0.14 µM (cyclic voltammetry) and 0.49 µM (amperometry) | Metrohm potentiostat | Water samples | Zambrano-Intriago et al. [12] |
| Enzymatic biosensor; colorimetric essay and chemiluminescence; photography followed by image analysis | Fusion of carbohydrate binding module from *Clostridium thermocellum* to a GP oxidase and a peroxidase; GP is oxidated to glyoxylate and $H_2O_2$ | Photography 0.25–2.5 mM; chemiluminescence reaction, 2–500 µM | Photography 0.12 mM; chemiluminescence reaction, 0.45 µM | Immobilization of enzymes to cotton buds, easy to hand and disposable assay | Water samples | Delprat et al. [13] |
| Enzymatic biosensor; electrochemical sensor | Inhibition of horseradish peroxidase by GP; a paper-based analytical device | 0.59–4.15 µM | 0.44 µM | Origami-like paper platform on three layers | Human urine | Moro et al. [14] |
| CHLDO·$FP_{phnG}$ WCB | Two different strains of *A. tumefaciens* transformed with pL1_$FP_{phnG}$ and were tested based on the double regulation of PhoB activator and phnF repressor TF | Fluorescence detection: 0.5–50 µM | 0.25 µM | Microplate readers with fluorescence measured at 620 nm (excitation at 530 nm); the naked eye for bacteria immobilized on paper filters | GBH samples, water samples, microbial cultures, soil treated with GBH | This work |

*Abbreviations*: N.R., not reported; FP, glyphosate; GBH, GP-based herbicides; TF, transcription factor; EPSPS, enzyme 5-enolpyruvylshikimate-3-phosphate synthase; WCB, whole-cell biosensor.

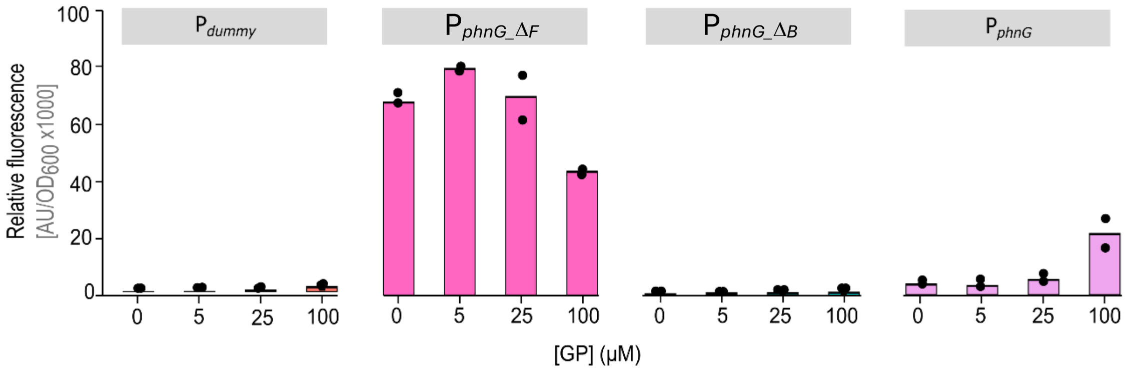


**Figure S1 · Performance of *A. tumefaciens* GV3101-based whole-cell biosensors upon incubation with increasing GP concentrations.** GV3101·$P_{dummy}$, GV3101·$P_{phnG\text{-}sf}$, GV3101·$P_{phnG\text{-}sb}$ and GV3101·$P_{phnG}$ cells were incubated in MSM with or without GP at different concentrations (indicated in the graphic, in µM) during 16 h. Maximum fluorescence (A.U./$OD_{600}$) values relative to the control, without GP are shown. Bars represent mean values ± standard deviation from three biological replicates. *A.U.*, arbitrary units; *$OD_{600}$*, optical density at 600 nm.

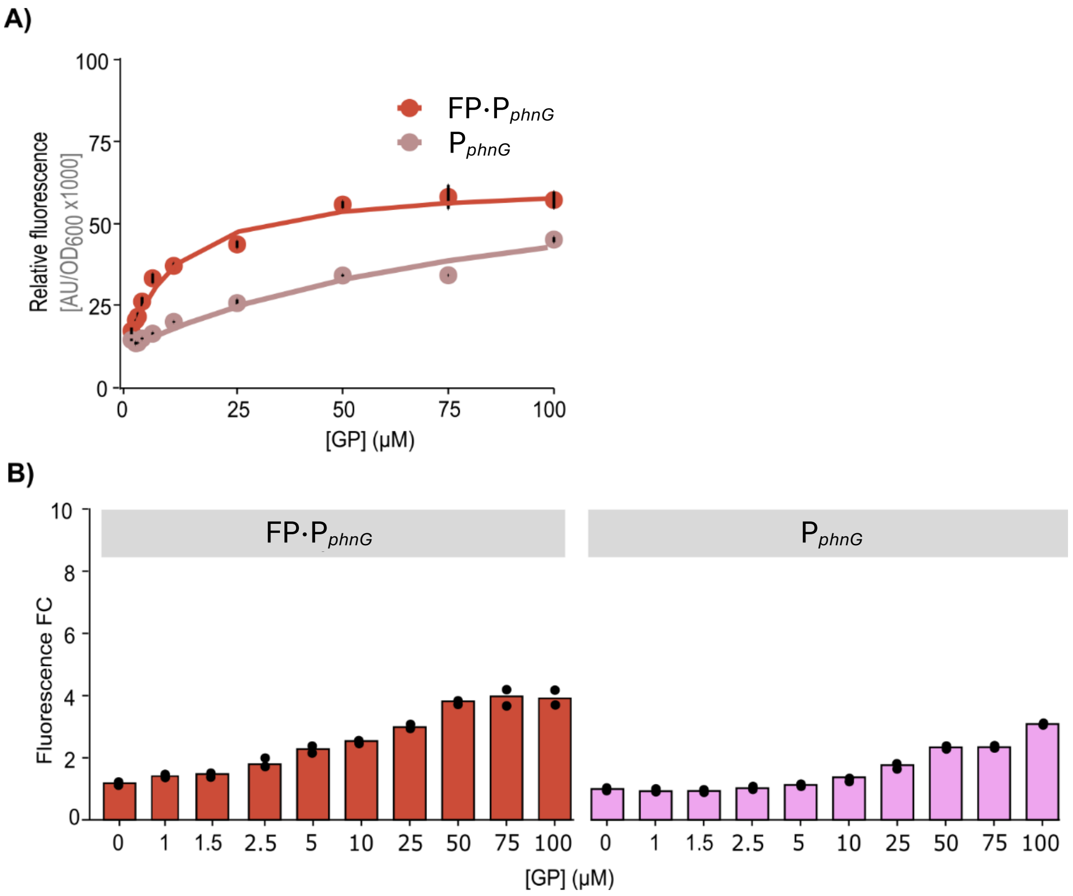


**Figure S2 · Performance of optimized *A. tumefaciens* GV3101-based whole-cell biosensors. (A)** GV3101·FP$_{phnG}$ and GV3101·P$_{phnG}$ were incubated in MSM with and without GP at different concentrations (1-150 µM) during 16 h. Dose-response trajectories for the optimized GV3101·FP$_{phnG}$ whole-cell biosensor (red) and the parental GV3101·P$_{phnG}$ biosensor (pink) are presented. **(B)** Maximum fluorescence (A.U./OD$_{600}$) values relative to the control, without GP are shown. Bars represent mean values ± standard deviation from three biological replicates. *A.U.*, arbitrary units; *OD$_{600}$*, optical density at 600 nm; FC, fold-change.

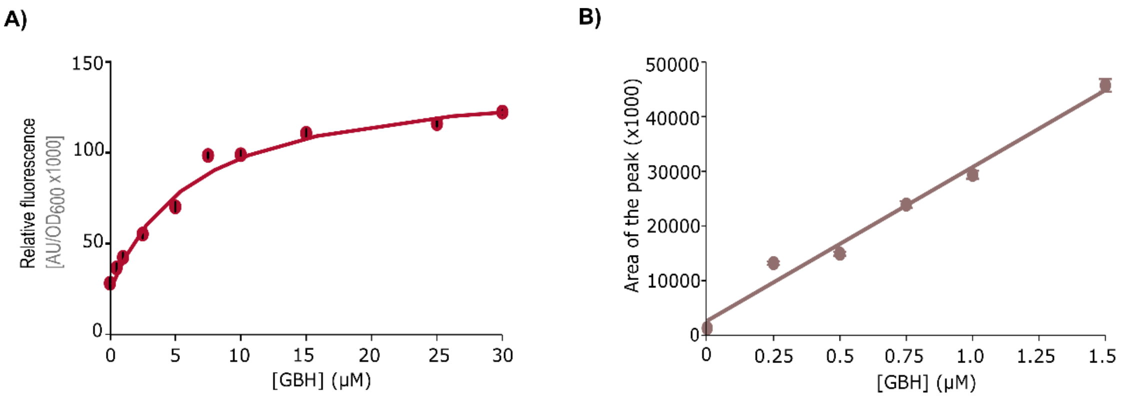


**Figure S3 · Response of the *A. tumefaciens* CHLDO·FP$_{phnG}$ whole-cell biosensor to GBH. (A)** Dose-response function for the CHLDO·FP$_{phnG}$ biosensor exposed to increasing GBH concentrations. **(B)** Fitting-curve for HPLC measurements of GBH. In both cases, results represent mean values ± standard deviation from three biological replicates. *A.U.*, arbitrary units; *$OD_{600}$*, optical density at 600 nm.
